# Mutation bias implicates RNA editing in a wide range of mammalian RNA viruses

**DOI:** 10.1101/2021.02.09.430395

**Authors:** Peter Simmonds, M. Azim Ansari

## Abstract

The rapid evolution of RNA viruses has been long considered to result from a combination of high copying error frequencies during RNA replication, short generation times and the consequent extensive fixation of neutral or adaptive changes over short periods. While both the identities and sites of mutations are typically modelled as being random, recent investigations of sequence diversity of SARS coronavirus 2 (SARS-CoV-2) have identified a preponderance of C->U transitions, potentially driven by an APOBEC-like RNA editing process. The current study investigated whether this phenomenon could be observed in the more genetically diverse datasets of other RNA viruses. Using a 5% divergence filter to infer directionality, 18 from 32 datasets of aligned coding region sequences from a diverse range of mammalian RNA viruses (including *Picornaviridae, Flaviviridae, Matonaviridae, Caliciviridae* and *Coronaviridae*) showed a >2-fold base composition normalised excess of C->U transitions compared to U->C (range 2.1x–7.5x). C->U transitions showed a favoured 5’ U upstream context consistent with previous analyses of APOBEC-mediated RNA targeting. Amongst several genomic compositional and structural parameters, the presence of genome scale RNA secondary structure (GORS) was associated with C->U/U->C transition asymmetries (*p* < 0.001), potentially reflecting the documented structure dependence of APOBEC-mediated RNA editing. Using the association index metric, C->U changes were specifically over-represented at phylogenetically uninformative sites, consistent with extensive homoplasy documented in SARS-CoV-2. Excess C->U substitutions accounted for 15-20% of standing sequence variability of HCV and other RNA viruses; RNA editing may therefore represent a potent driver of RNA virus sequence diversification and longer term evolution.

**Author Summary:** The rapid evolution of RNA viruses is thought to arise from high mutation frequencies during replication and the rapid accumulation of genetic changes over time in response to its changing environments. This study describes an additional potent factor that contributes to the evolution of RNA infecting mammals, the deliberate mutation of the viral genome by host antiviral pathways active within the cell when it becomes infected. This so called “genome editing” by one or more APOBEC enzymes leads to characteristic C->U mutations that damage the virus’s ability to replicate. While this pathway is well characterised as an antiviral defence against HIV and other retroviruses, this study provides evidence for its activity against a wide range of human and veterinary viruses, including HCV and foot and mouth disease virus. APOBEC-driven mutations accounted for 15-20% of standing sequence variability of RNA virus groups, representing a potent driver of RNA virus sequence diversification.

## INTRODUCTION

The evolution of viruses is typically conceptualised as a combination of adaptive sequence change in response to a range of selection pressures in the environment and a process of random diversification in which neutral or near neutral nucleotide substitutions become fixed in virus populations (1-3). An extensive literature documents adaptive (or Darwinian) evolution in response to antiviral treatments and escape from host immune responses in the form of T cell and antibody driven epitope escape mutation (4-7). Viruses may furthermore display a series of changes to encoded viral proteins and even genome rearrangements in the process of jumping hosts and adapting to new internal environments (8, 9).

A further source of mutations in viruses arises from effects of several innate antiviral effector mechanisms in vertebrate cells that operate through viral genome editing. Of these, the best characterised are the interferon-inducible isoform of adenosine deaminase acting on RNA type 1 (ADAR1)(10) that targets RNA viruses during replication, and members of the apolipoprotein B mRNA-editing enzyme, catalytic polypeptide-like (APOBEC) family (11). The substrate for ADAR1 is double-stranded (ds) RNA formed as a replicative intermediate; upon binding, it catalyses the deamination of adenine bases to inosine which are subsequently copied as a G by viral RNA polymerases, creating A->G base mutations (or U->C on the opposite strand). Members of the APOBEC family typically target single stranded DNA or RNA templates for mutagenesis of cytidine to thymidine. For example, APOBEC3G edits the single stranded DNA generated by reverse transcription of retroviruses. The resulting mutated proviral copy is unable to direct further retroviral replication (12-14). Consistent with its role in viral defence, the APOBEC3 gene locus shows evidence for rapid expansion through gene duplication in mammals, with the human genome encoding seven active antiviral proteins (A3A, A3B, A3C, A3D, A3F, A3G and A3H). As a measure of the effectiveness of this pathway, exogenous retroviruses invariably encode antagonists of APOBEC3, such as Vif in HIV-1, to promote its degradation by ubiquination (15) or prevent its incorporation into virions (16).

While these viral DNA and RNA editing pathways have been recognised and functionally investigated for over two decades, whether they make a substantial contribution to the longer term evolution of viruses remains unknown, while their mutagenesing effects on the generation of viral diversity remain outside of mainstream evolutionary models. Both the phenomenon of G->A hypermutation in HIV-1 (17) through the actions of APOBEC 3G (13, 14) and ADAR1-mediated editing of measles virus genomes associated with sub-acute sclerosing panencephalitis (18) are readily observable but both are effectively evolutionary dead-ends and do not contribute to the ongoing pool of replicating virus lineages. It had indeed been proposed that Vif in HIV-1 is so effective in counteracting effects of APOBEC that it prevents any genomic signatures of previous editing in the HIV genome (19). Indeed, one of the difficulties with observing the occurrence of editing in RNA viruses is the likelihood that one or a small number of RNA edits will render modified virus substantially less fit or entirely replication incompetent and therefore unrepresented in naturally occurring virus populations. Effects of editing may therefore only be manifest at a small minority of previously unedited sites where mutations have minimal effects on viral fitness. However, the recent assembly of large numbers of accurate full genome sequences of SARS-CoV-2, and minimal naturally occurring sequence variability (20) has provided a rare opportunity to observe editing effects on a relatively blank genomic canvas. In such sequences, we and others have reported the occurrence of a substantial excess of C->U transitions among genomic sequences (21-24), in one study shown to represent over 40% of all mutations observable in currently compiled sequence datasets (21). Despite the relative abundance of C->U transitions and their contribution to the overall substitution rate of the virus, their effect on the longer term evolution of SARS-CoV-2 may be limited. C->U mutations are typically homoplastic and revert to the original wild type cytidine in descendant lineages (21, 22), as if representing transiently tolerated less fit mutants that are eliminated from the virus population.

Although hitherto generally considered as an antiviral pathway primarily active against retroviruses and retroelements, the evidence for extensive APOBEC-mediated editing of coronavirus genomic RNA sequences naturally leads to the question of whether such processes may operate on other RNA viruses. Evidence for such editing may include observations of an excess numbers of C->U changes over U->C changes where directionality can be inferred. Suspected edited sites may additionally possess the same 5’ or 3’ base contexts that are known to favour APOBEC-mediated editing of retroviral genomes.

In the current study, alignments of mammalian RNA virus sequences have been assembled from a range of families for which multiple full genome sequences have been obtained. Since APOBEC-mediated RNA editing has been proposed to occur in the context of RNA structure elements, virus datasets were additionally scanned for sequence-dependent internal RNA base-pairing (25, 26). Virus genome structural and compositional factors favouring APOBEC-like C->U editing in these datasets were analysed and the potential contribution of these driven sequence changes to their longer-term evolutionary trajectories estimated.

## RESULTS

### Sequence datasets

The primary resource for the study were aligned sequences from a wide range of mammalian RNA viruses derived from several vertebrate virus families (Table 1; Table S1; Suppl. Data). These were selected based on the availability of large numbers of complete genome sequences from naturally occurring virus variants collected in previous epidemiological and evolutionary studies. These showed levels of intra-population sequence divergence ranging from 5%-19% (Table 1). These included viruses with and without large scale RNA secondary structure in the genomes (25-27), with mean folding energy differences (MFEDs) ranging from -0.1% - 17.8% (Table 1). The analysis was supplemented by inclusion of previously described datasets of SARS-CoV-2 and other coronaviruses (27).

**TABLE 1.**
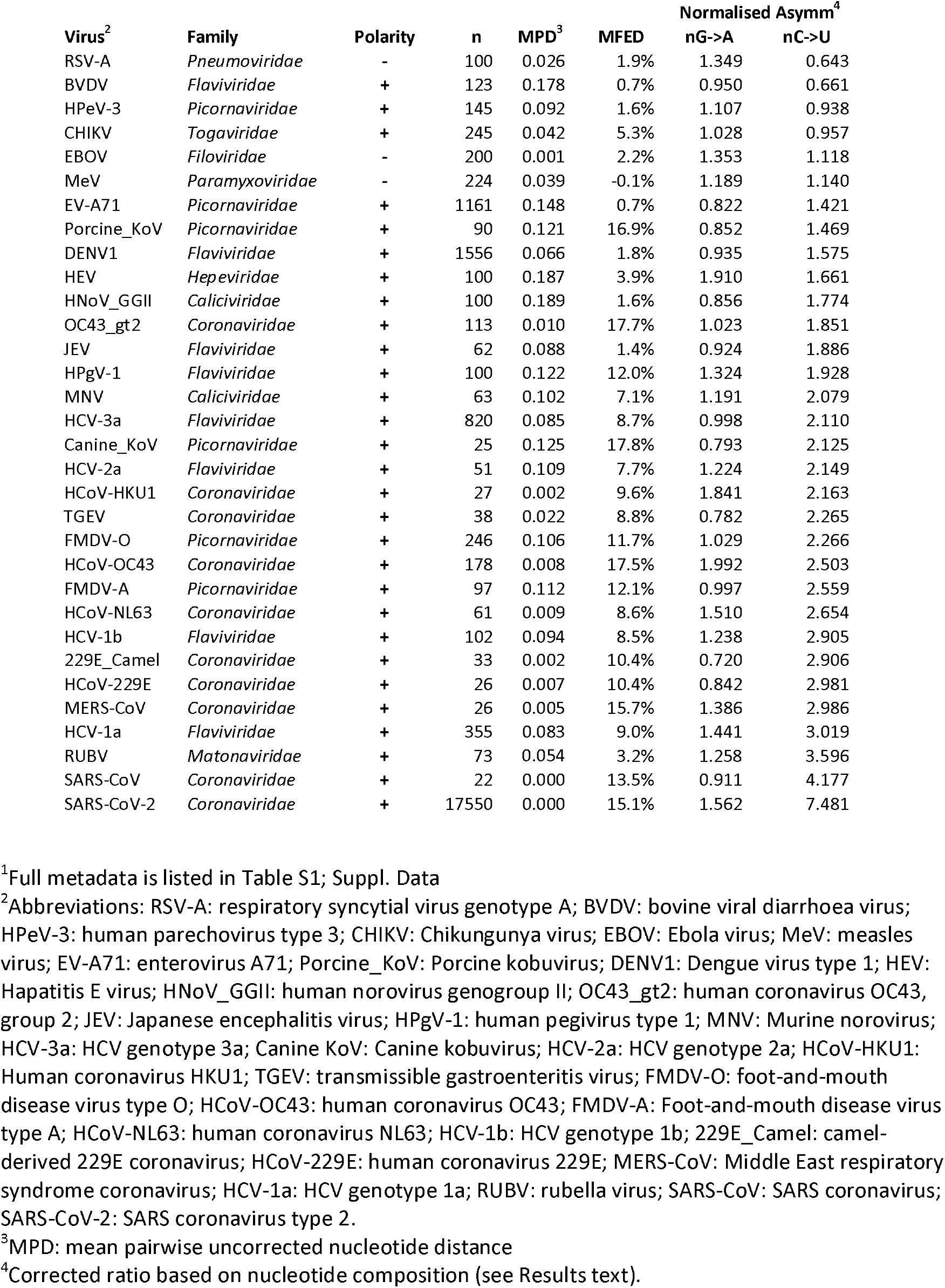
COMPOSITIONAL FEATURES OF RNA VIRUS SEQUENCE DATASETS USED IN THE STUDY^1^

### Detection of mutational asymmetries in RNA virus datasets

The previous analysis of the directionality of sequence changes in SARS-CoV-2 and the detection of an excess number of C->U changes was simplified by the minimal sequence diversity of the assembled post-pandemic sequences (20). SARS-CoV-2 sequence diversity primarily comprised isolated base changes relative to a consensus sequence shared by all but one or a few sequences in the alignment. However, for the datasets analysed in the current study, population diversity was substantially greater making inference of directionality increasingly arbitrary as site variability increased. To overcome this problem, analyses of relative mutation frequencies were restricted to sites showing low degrees of heterogeneity so that the directionality of mutations can be inferred.

Sequence datasets was analysed using the program Sequence Change in the SSE package, which records the occurrences and sites of sequence changes from a majority rule alignment consensus sequence. Collectively, there was a significantly greater number of C->U changes compared to other transitions in the RNA virus datasets at sites showing <5% heterogeneity although the degree of over-representation was highly variable between viruses (Fig. 1A). To more formally quantify the degree of over-representation of C->U transitions in each virus dataset, the ratio of each transition to its reverse (Columns 13; 14, Table S1; Suppl. Data) was normalised for base composition as described in a previous analysis for SARS-CoV-2 (21) (Columns 7, 8, Table 1). Formally, in the absence of mutational pressure (null expectation), the expected ratio of frequencies of a mutation X->Y to Y->X would be proportional to their native base frequencies and can be normalised as [f(X->Y) / f(Y->X)] / f(Y) * f(X). Applying this to the observational data in the virus datasets, the high frequencies of C->U changes were reflected in strong C->U / U->C transition asymmetries (Fig. 1B; mean value 2.3), again though with a wide range of values from 0.7 (BVDV) – 7.5 (SARS-CoV-2). Contrastingly, the complementary G->A / A->G transition values were less variable and centred around the null expectation (mean 1.2, range 0.6 -2.4).

**FIGURE 1.**
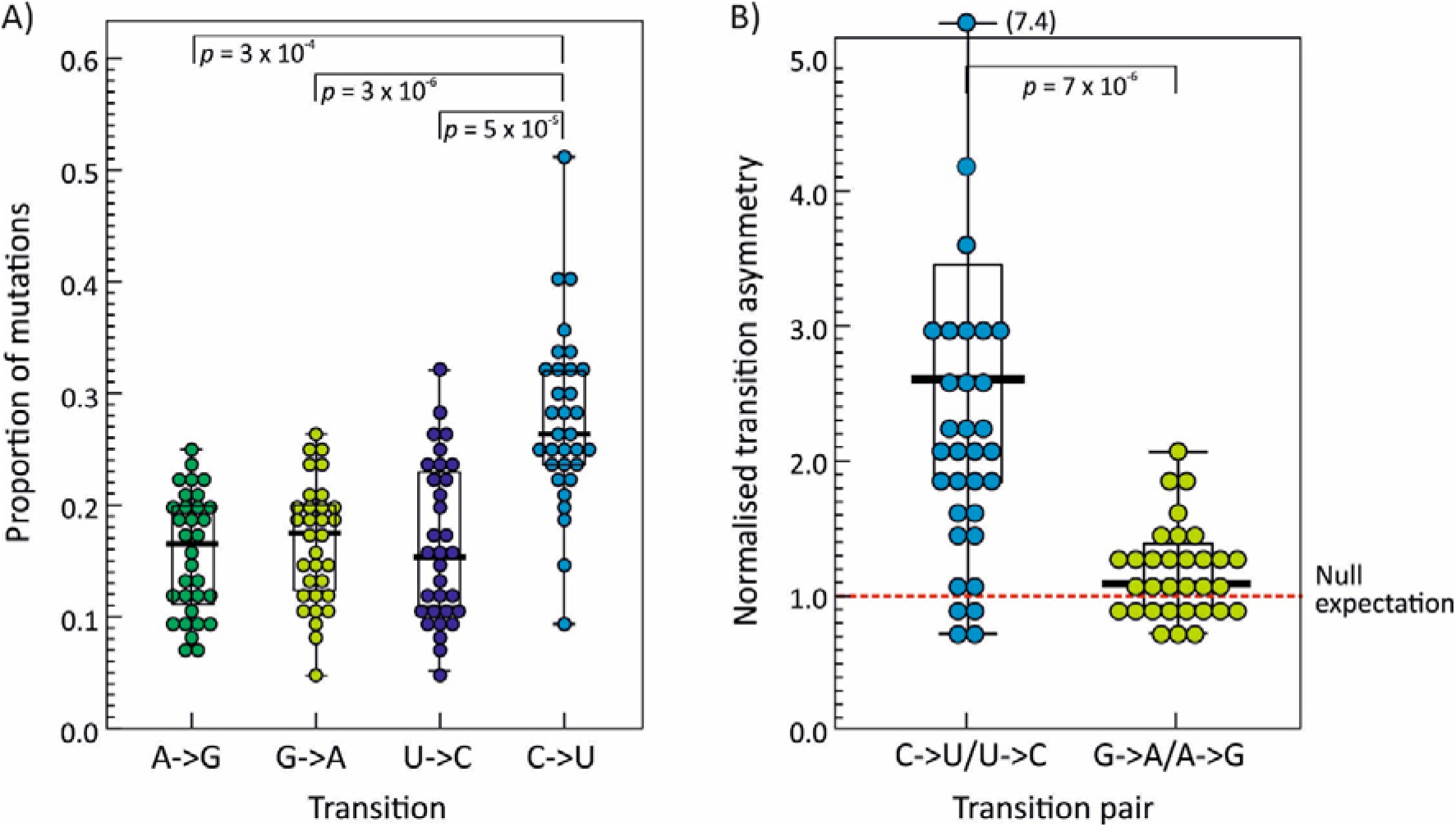
TRANSITION FREQUENCIES AND ASYMMETRIES IN RNA VIRUS ALIGNMENTS. (A) Relative frequencies of each mutation type expressed as a percentage of all changes (y-axis) in the 32 RNA virus alignments at sites showing <5% heterogeneity. (B) Comparison of normalised transition asymmetry values; the dotted red line shows the expected unbiased transition asymmetry. For both graphs, distributions were compared using the Friedman’s two-way analysis of variance by ranks test (with Bonferroni correction for multiple comparisons in Fig. 1A); only significant (*p* < 0.05) two-tailed *p* values shown. Box plots show maximum, upper interquartile range (IQR), median, lower IQR and minimum values of each distribution.

This initial analysis of mutational asymmetry was conducted using a 5% heterogeneity threshold to allow directionality of sequences to be inferred. The relationship between site heterogeneity and transition asymmetry was determined for two example RNA virus datasets showing excess C->U changes at the 5% threshold (Fig. 2; HCV – mean transition asymmetry 3.0; FMDV: 2.2; Table 1). At highly variable sites, frequencies of G->A / A->G and C->U / U->C were comparable and close to the null expectation. However, for both viruses, increasing asymmetry was observed at sites with reduced heterogeneity, ruling out the possibility that the observed asymmetries were the result of unrecognised compositional biases in the virus datasets, and validating the use of the 5% threshold to analyse directionality of sequence change. Similar relations were observed for other viruses showing elevated C->U / U->C transition asymmetries (data not shown).

**FIGURE 2.**
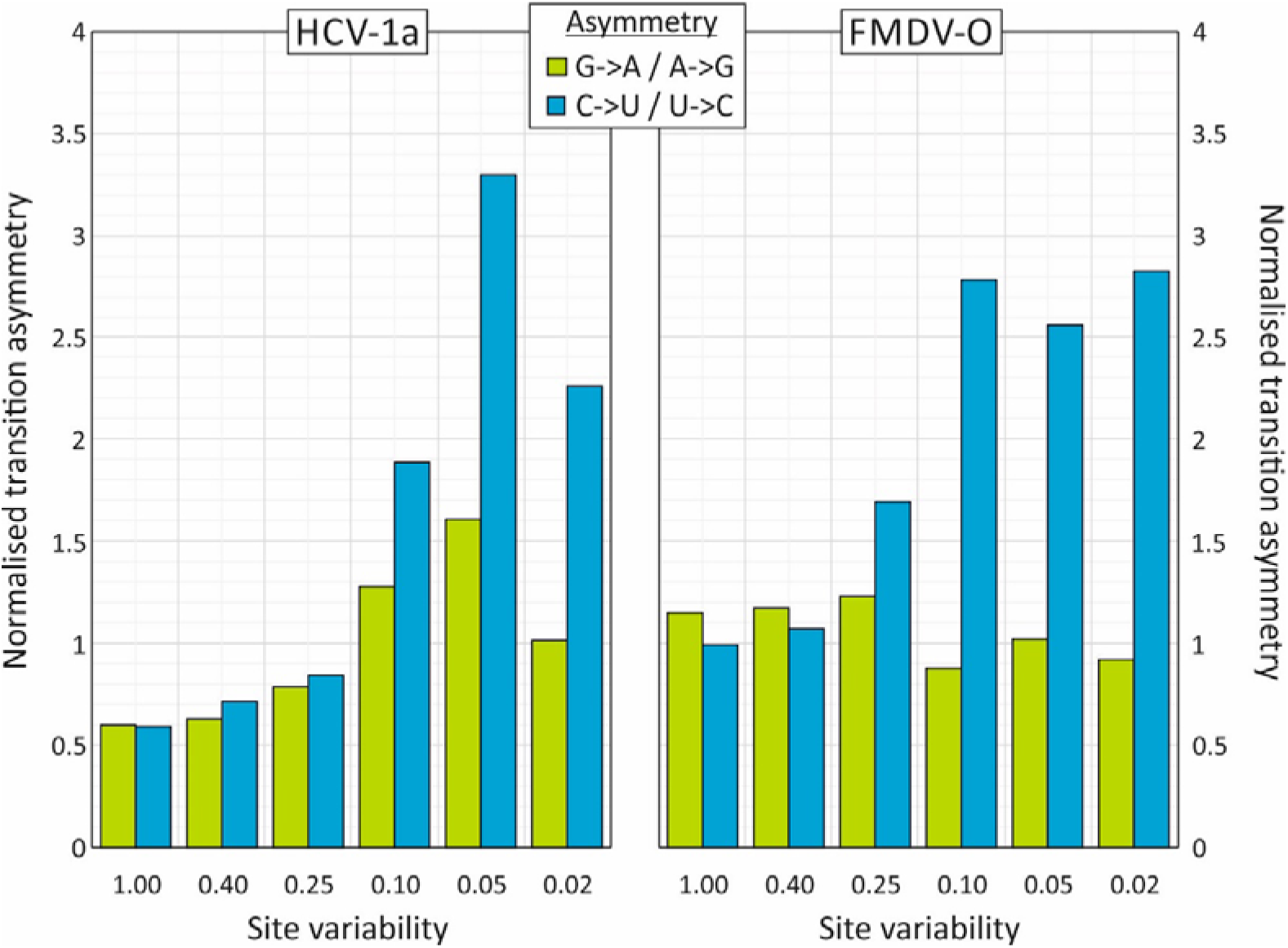
FREQUENCY RELATED TRANSITION ASYMMETERIES. Transitional asymmetries of two virus datasets showing C->U/ G->A asymmetries. Normalised values (y-axis) were calculated for sites showing different levels of sequence heterogeneity (x-axis): 0.02: 0.02 or less; 0.05: <0.05 and ≥0.02; 0.1: <0.1 and ≥0.05; 0.25: <0.25 and ≥0.1; 0.4: <0.4 and ≥0.25; 1.00: ≥0.4.

The estimation of relative transition frequencies was collectively based upon all sequences within each virus alignments. To investigate the degree of heterogeneity in C->U changes between sequences, numbers of this transition were computed individually and compared with those of the reverse mutation (Fig. 3) in two of the larger datasets showing high and low normalised transition asymmetries (HCV-1a – 3.1 and EV-A71 – 1.4 respectively; Table 1). For EV-A71, there were means of 3.1 C->U and 2.5 U->C substitutions per sequence at the 5% heterogeneity level, and a distribution of values that approximated to a Poisson distribution, although marginally over-dispersed (Kolmogorov-Smirnov single sample test statistic = 6.8; *p* < 0.001). Similarly for HCV-1a, both U->C and C->U transition frequencies followed marginally skewed Poisson distributions (test statistics 1.5 [*p* = 0.03] and 2.5 [*p* < 0.001] respectively), but with a higher mean number of C->U transitions per sequence (12.1 / sequence) than the reverse (2.8 / sequence). The sequence datasets of HCV and EV-A711 sequences therefore showed no evidence for the occurrence of individual hypermutated sequences as described previously for HIV-1 (17, 28, 29); the driver of elevated C->U frequencies appeared to operate at a similar intensity on all sequences analysed in each dataset.

**FIGURE 3.**
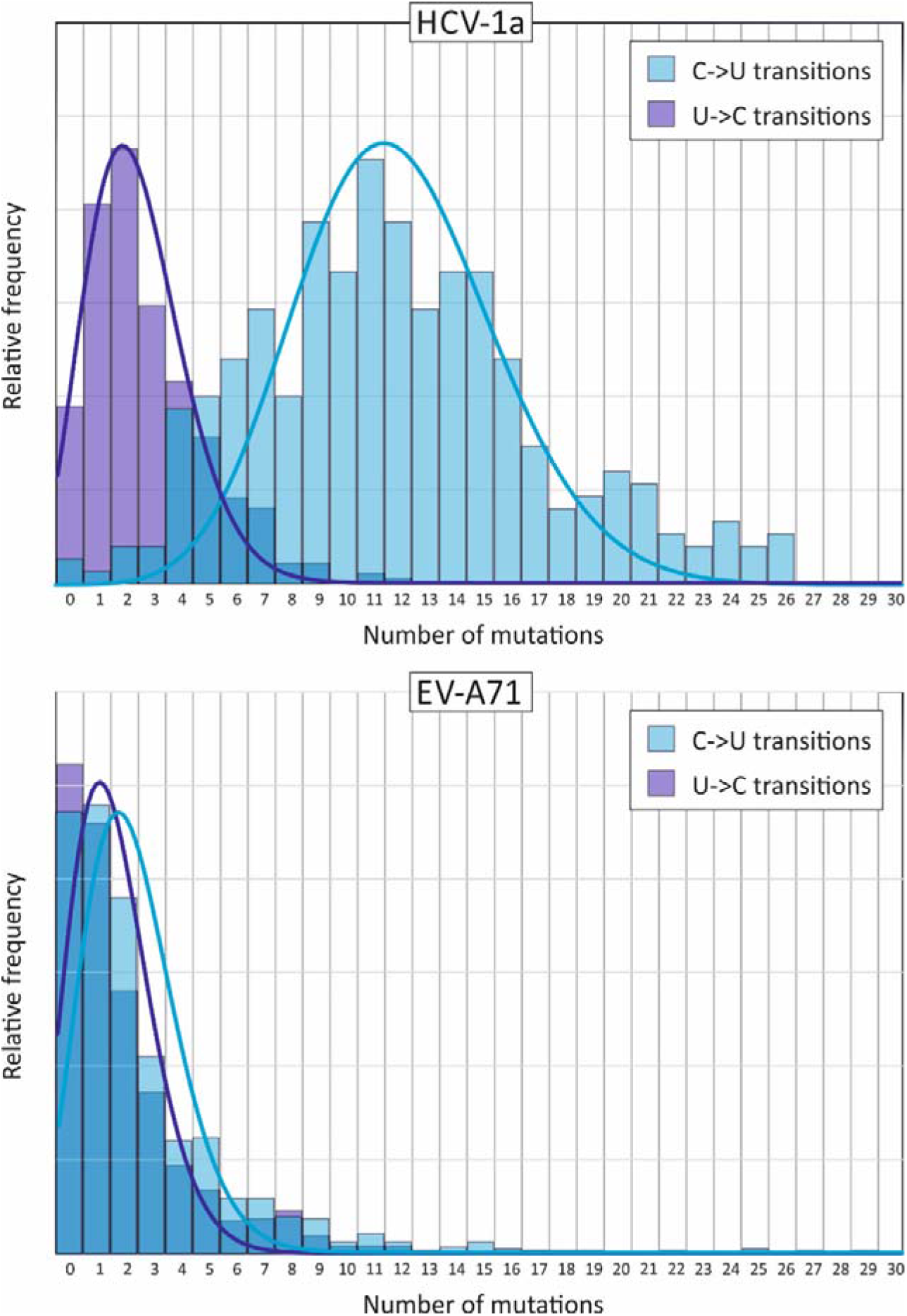
DISTRIBUTION OF C->U AND U->C MUTATIONS IN INDIVIDUAL SEQUENCES. Numbers of C->U and U->C transitions in individual coding region sequences of HCV-1a and EV-A71 plotted as frequency histograms. Distributions were fitted to Poisson distributions based around their mean numbers of substitutions (light and dark blue lines).

To investigate which genome features of RNA genomes were predictive of the C->U/U->C transition asymmetry, a range of compositional attributes (G+C content, representation of CpG and UpA dinucleotides, asymmetry in the number of G bases relative to C, and of A relative to U), mean folding energies (MFEs) of consecutive 300 base fragments and differences of this value from sequence order randomised controls (MFEDs) were computed. The association of each with C->U/U->C and G->A/A->G transition asymmetry values was analysed by multivariate analysis (Table 2). MFED value was the only variable significantly associated with C->U / U->C asymmetry (p = 0.045); this parameter represents the degree of sequence order-dependent RNA folding in the coding region(s) of the virus (Table 2). The association between C->U / U->C asymmetry and MFED was further apparent and readily visualised by simple linear regression (Fig. 4; *p* = 0.005). There was no association with MFE, representing the minimum free energy on RNA folding, a property primarily influenced by the G+C content of the sequence, which also showed no association with the C->U / U- >C transition asymmetry. There were similarly no associations with extents of CpG or UpA dinucleotide suppression in RNA virus genomes, or base imbalances (U/A, C/G). As expected from the minimal differences from the null expectation, no association between G->A/A->G transition asymmetry values with any compositional or structural sequence attribute was detected.

**TABLE 2.**
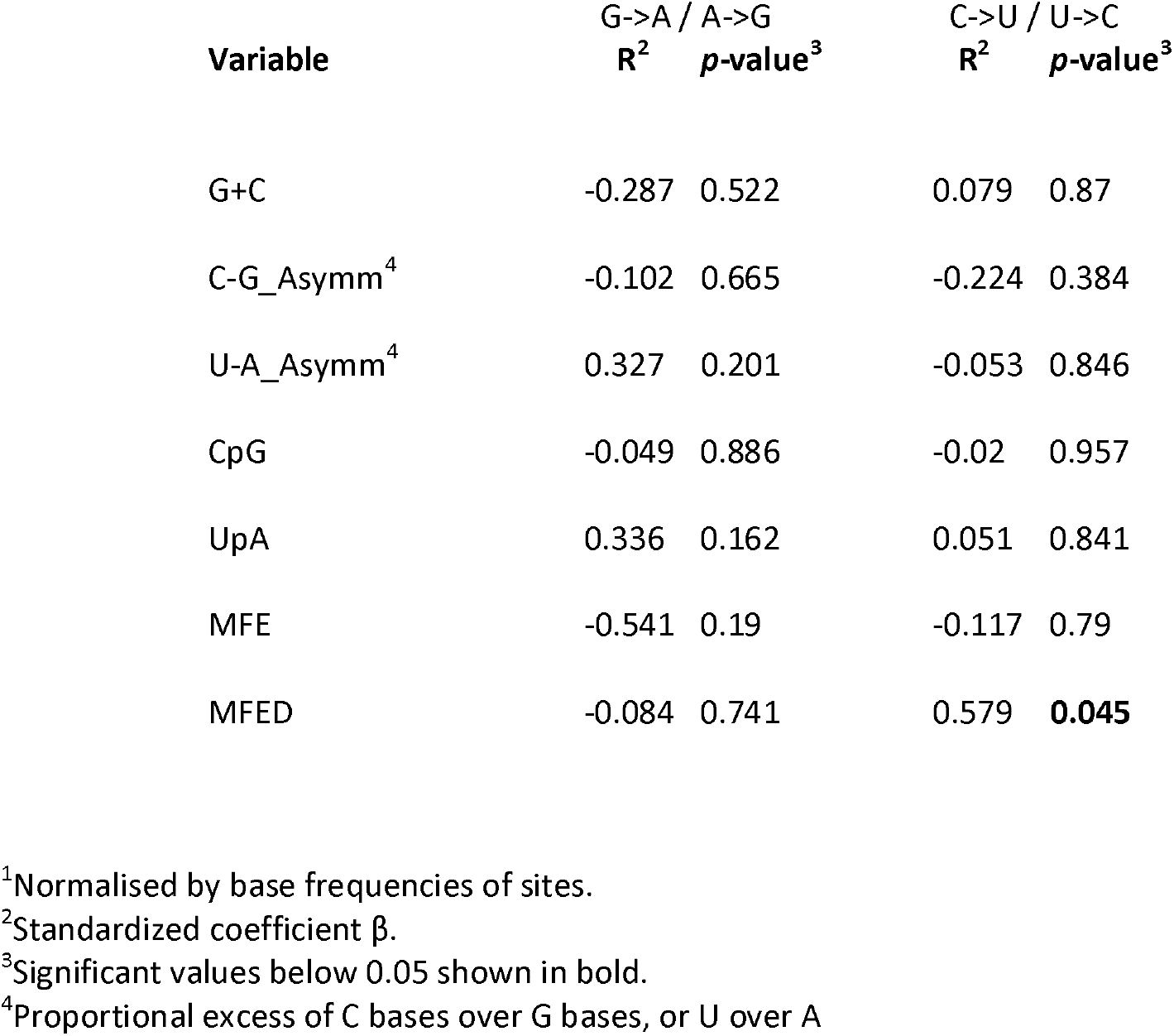
PREDICTIVE FACTORS FOR G->A AND C->U TRANSITION ASYMMETRIES^1^ BY ANOVA

| Variable | G->A / A->G |  | C->U / U->C |  |
| --- | --- | --- | --- | --- |
|  | R<sup>2</sup> | p-value<sup>3</sup> | R<sup>2</sup> | p-value<sup>3</sup> |
| G+C | -0.287 | 0.522 | 0.079 | 0.87 |
| C-G_Asymm<sup>4</sup> | -0.102 | 0.665 | -0.224 | 0.384 |
| U-A_Asymm<sup>4</sup> | 0.327 | 0.201 | -0.053 | 0.846 |
| CpG | -0.049 | 0.886 | -0.02 | 0.957 |
| UpA | 0.336 | 0.162 | 0.051 | 0.841 |
| MFE | -0.541 | 0.19 | -0.117 | 0.79 |
| MFED | -0.084 | 0.741 | 0.579 | **0.045** |

<sup>1</sup>Normalised by base frequencies of sites.
<sup>2</sup>Standardized coefficient $\beta$ .
<sup>3</sup>Significant values below 0.05 shown in bold.
<sup>4</sup>Proportional excess of C bases over G bases, or U over A

**FIGURE 4.**
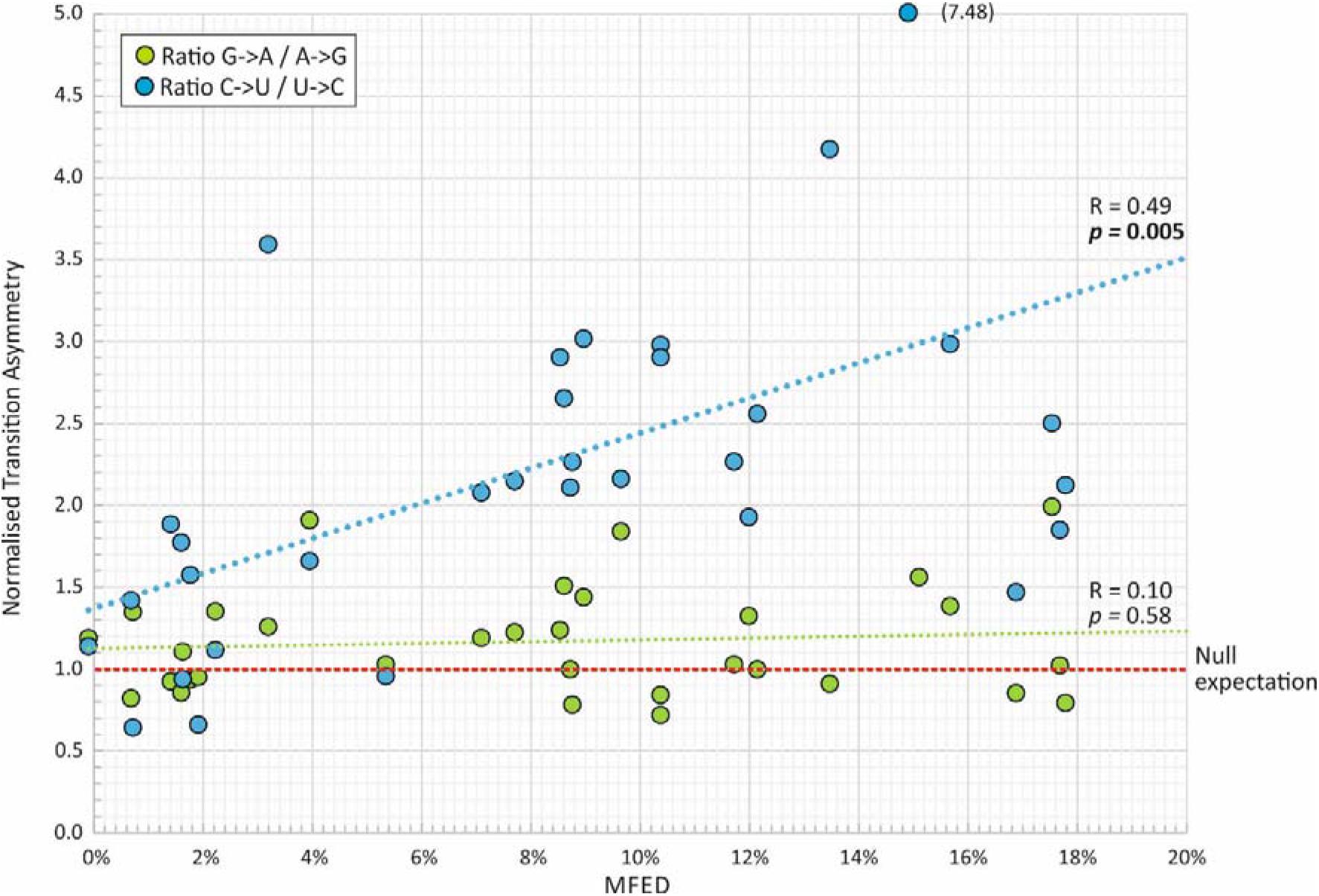
ASSOCIATION OF TRANSITION ASYMMETRIES WITH RNA SECONDARY STRUCTURE. The association of transition asymmetry values with MFED values, indicating of the degree of genome RNA folding. Correlation values (R) and significance using linear regression for C->U / U->C and G->A / A->G asymmetries are shown.

### Sequence contexts for C->U transition asymmetry

C->U transitions in SARS-CoV-2 genomes were influenced by the immediate 5’ and 3’ base contexts of the mutated site (21). Other RNA virus datasets showing C->U / U->C transition asymmetries were analysed similarly (Fig. 5). Once normalised to base composition, relative mutation frequencies varied over a substantial range but with a 5’U being consistently associated with greater C->U/U-C transitional asymmetry. Sites with a 5’U showed a mean over-representation for all viruses of 2.0 compared to 0.78, 0,56 and 0.,84 in 5’ A, C and G contexts respectively (*p =* 0.0004, 5 x 10 and 8 x 10-5 respectively by Kruskall Wallace non-parametric test). Effects of 3’ context were far more variable between viruses, but with evidence for favoured C->U transitions upstream of A in FMDV and to lesser extents in other RNA viruses.

**FIGURE 5.**
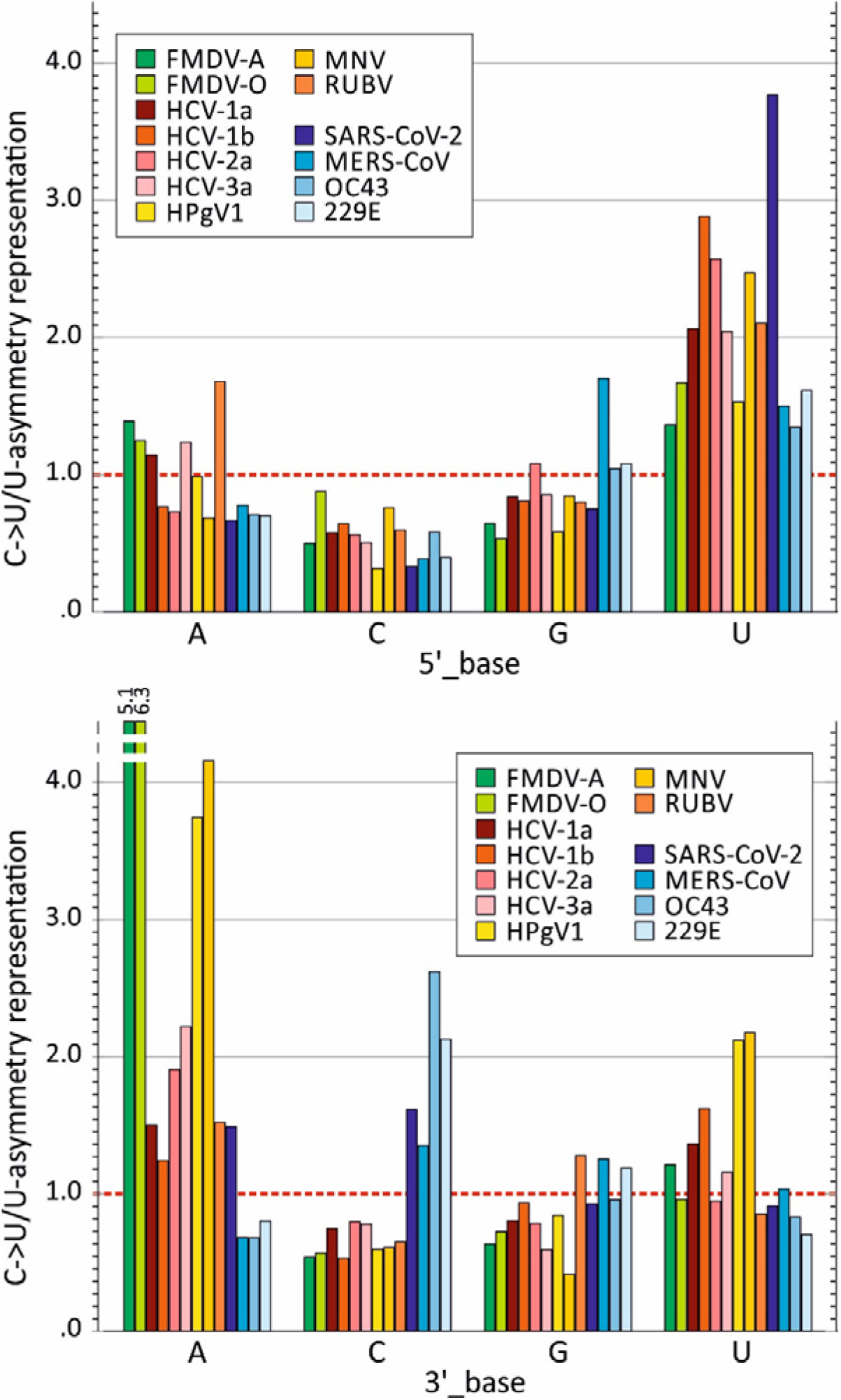
INFLUENCE OF 5’ AND 3’ BASES ON C->U MUTATION FREQUENCIES. Influence of the identities of the immediate 5’ base and 3’ base on C->U mutation frequencies in a range of RNA viruses showing C->U/U->C transition asymmetry. Normalised C->U/C->U transition asymmetries in each 5’ and 3’ context were adjusted to account for 5’ or 3’ base frequencies. The y-axis shows the over- or under-representation of the asymmetry values in each context relative to the value for all contexts; the null expectation (no effect of 5’ or 3’ base) was 1.0 (red dotted line).

### Homoplasy of C->U mutations

Previous analyses demonstrated that a proportion of C->U mutations in SARS-CoV-2 failed to become genetically fixed in a population (22). The distribution of many mutations violated the overall phylogeny of the dataset, appearing convergently and transiently in different parts of the tree (21). The possibility that excess C->U changes observed in current datasets might be similarly homoplastic was investigated more systematically through measurement of the concordance between nucleotide identities at variable sites in virus alignments and their overall tree topology. This enables segregating substitutions that reflect evolutionary relationships to be distinguished from phylogenetically uninformative or incongruent sites that may arise from host-driven mutational processes. As the method does not require directionality to be inferred, the approach is not restricted to relatively invariant sites (<5% heterogeneity) examined in previous analyses.

The program, Homoplasy Scan in the SSE package was developed to sequentially analyse each variable site in a virus sequence alignment; this recorded the degree of segregation of each base in a global tree constructed from 1200 base genome fragment that incorporates the interrogated bases (Fig. 6). Association index (AI) values based on sequences grouped by their component bases were typically low in DENV3, HPeV-3 and EV-A71, indicating that most substitutions co-segregated with overall phylogeny. AI distributions were typically narrower (more informative) for more variable sites (high Shannon entropy values) despite the potential effect of site saturation and convergence on site with only 4 possible character states. The pattern of base segregation was remarkably different in alignments of HCV (results from genotype 1a are shown but other genotypes were closely similar; data not shown) and HPgV-1 (Fig. 7). In these, only a small fraction of sites showed a base distribution that segregated with (and defined) the overall phylogeny of the alignment; the majority, irrespective of their underlying diversity, poorly matched overall phylogeny with AI values approaching the mean of the null distribution (AI = 1.0). Distributions for FMDV, MNV and JEV were intermediate between these extremes.

**FIGURE 6.**
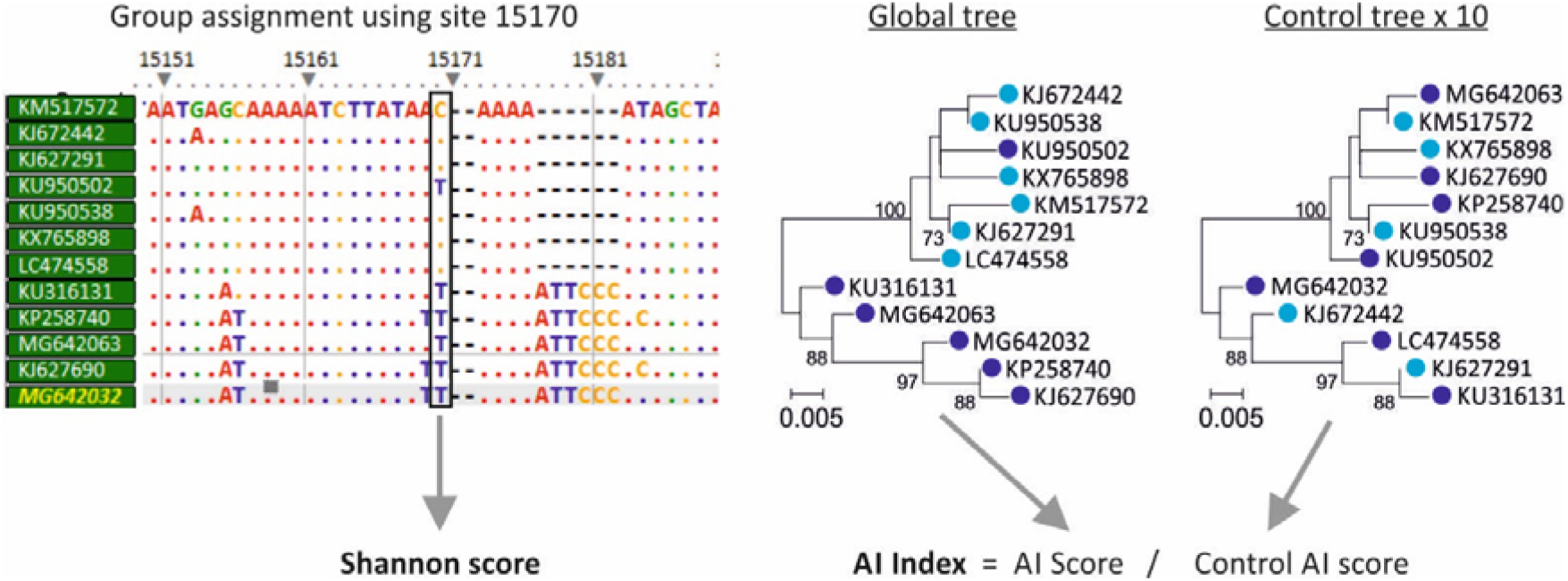
USING THE ASSOCIATION INDEX TO DETERMINE INFORMATIVNESS OF INDIVIDUAL SITES. Schematic summary of the steps used to investigate site informativeness. Individual alignment positions are sequentially analysed for their concordance to a global phylogeny. Base identity is used to assign groups which are then use for calculation of an association value through group segregation in a neighbour-joining tree of the alignment where non-bootstrap supported branches are collapsed. The AI index is its ratio to the mean association value of 10 sequence label order randomised controls (representing the null expectation of no association). Finally, the Shannon entropy score, representing site heterogeneity is recorded.

**FIGURE 7.**
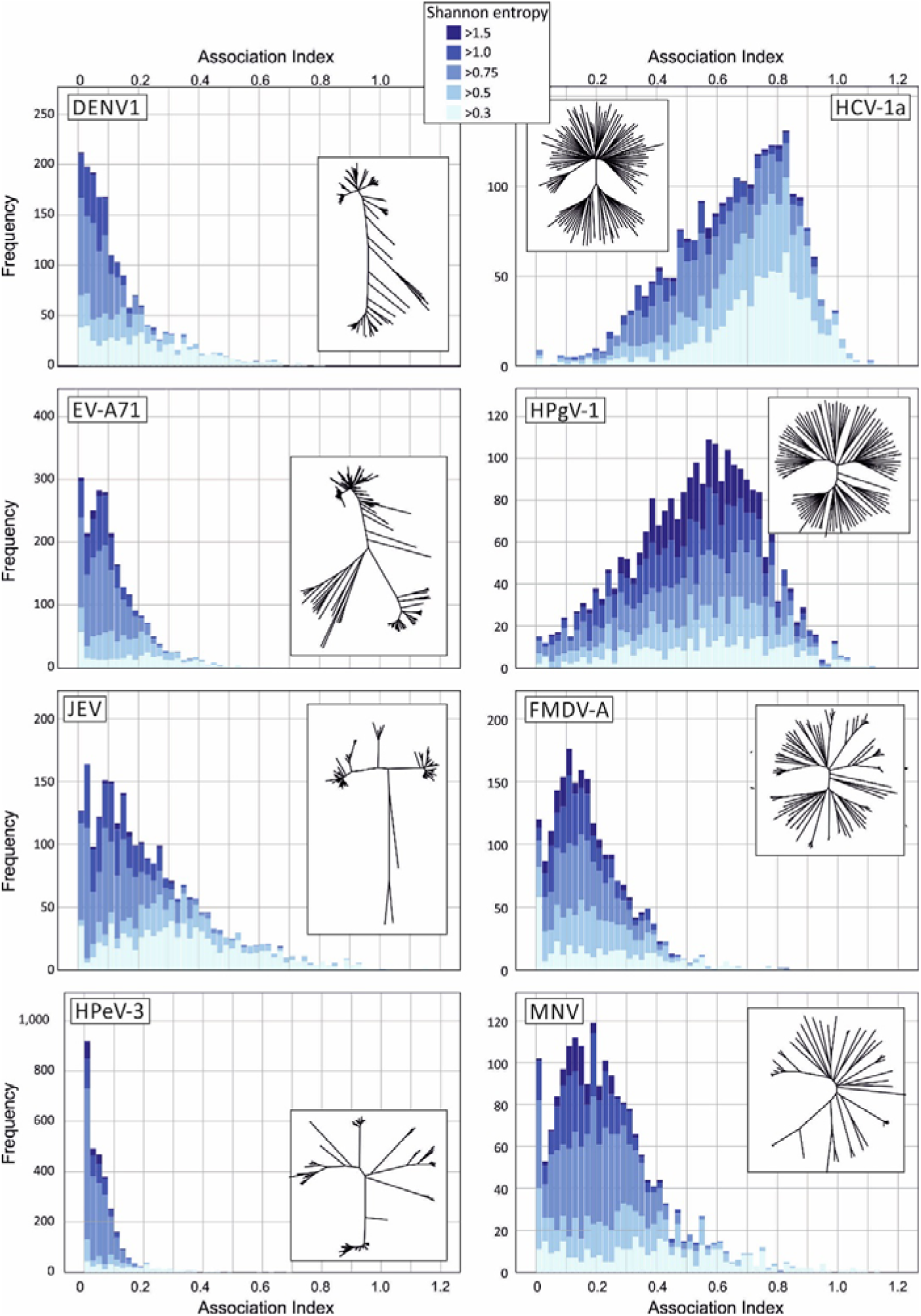
DISTRIBUTION OF ASSOCIATION INDEX VALUES IN VIRUS DATASETS. Frequency distributions of AI values at variable sites in alignments of representative viruses showing unbiased (left) or elevated (right) C->U/U->C transition asymmetries and with comparable overall sequence divergence (MPD values listed in Table 1). Histograms were sub-divided based on their Shannon entropy range (see key for colour coding; minimally variable sites (Shannon entropy < 0.3) were excluded). Insets show the corresponding tree topologies for each virus analysed, for large datasets (HCV, EV-A71; DENV1), trees based on randomly selected representative sequences are shown for clarity. Phylogenetic trees drawn to scale are provided in Fig. S1 (Suppl. Data).

Broadly, the observed differences arose from different genetic structuring of sample populations; phylogenetic trees from alignments with predominantly informative sites showed a marked degree of structured internal branching (Fig. S1; Suppl. Data), while variants of HCV and HPgV-1 showed deep branching and little ordering of sequences beyond their initial diversification. The differences in tree structure between datasets were quantified using a lineage through time plot generated by the LTT program in the Phylocom package (30) (Fig. S2; Suppl. Data). This depicts the substantial lineage diversification of HCV, HPgV-1 and to lesser extents of MNV and FMDV-A at the base of the tree, and a contrasting late diversification of JEV, DENV1, HPeV-3 and EV-A71.

Irrespective of the actual distributions of AI values (and associated differences in tree topologies of the different RNA virus datasets), categorisation of sites based on AI value ranges allowed an investigation of whether C->U transition frequencies were specifically over-represented at phylogenetically uninformative sites as would be expected if these mutation were subject to homoplasy (Fig. 8). Amongst representative viruses showing C->U/U->C transition asymmetry (HCV-1a, HPgV-1, MNV and FMDV-O), there was a significant excess of C->U transitions compared to other transitions at uninformative sites (AI values > 0.2; Fig. 8; upper left graph), in contrast to ratios observed in the unbiased dataset (EV-A71; JEV, DENV1 and HPeV-3; upper right graph). Excess C->U transitions were also more extensively distributed at sites of medium / low Shannon entropy values (lower left graph). Contrastingly, the representation of other transitions (U->C, G->A and A->G) showed no association with either AI score or site variability. The relationship between excess C->U changes at sites with high AI values and low variability is consistent with the hypothesis for extensive homoplasy specifically of this transition.

**FIGURE 8.**
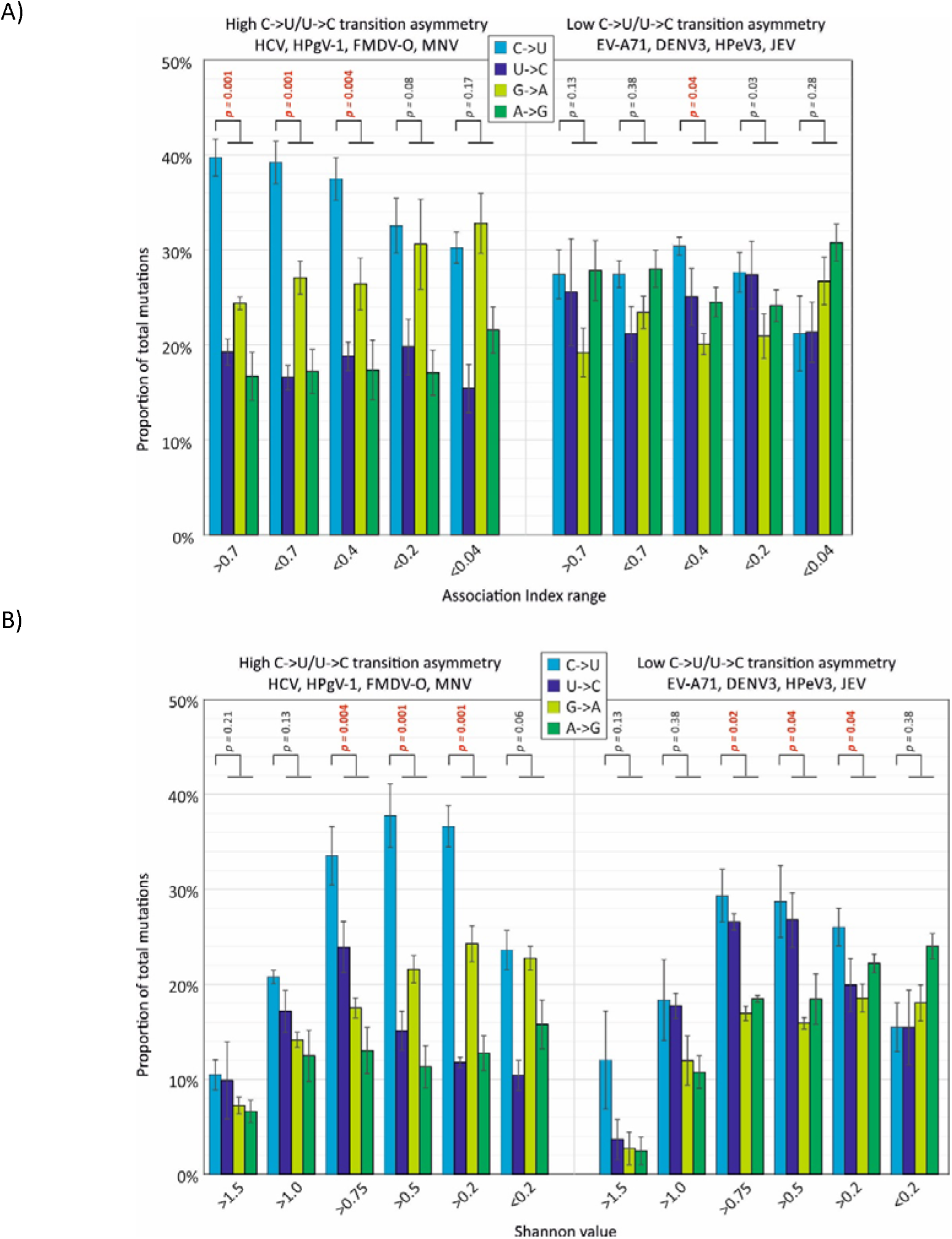
EFFECT OF ASSOCIATION INDEX VALUES AND SITE VARIABILITY TRANSITION FREQUENCIES. Relative frequencies of different transitions at sites varying in AI value, reflecting their phylogenetic informativeness (A), and in sequence heterogeneity (B). Bar heights show means of the four component virus datasets; error bars show standard errors of the mean). Frequencies of C->U were compared with frequencies of the other three transitions in each band using the Mann-Whitney U test; significant values shown in red.

### Contribution of C->U transitions to viral diversity

The extent to which the observed excess of potentially homoplastic and transient C->U mutations in certain viral datasets contributed to overall viral sequence diversity was calculated. The number of sites in a virus alignment showing a majority C->U change subtracted by those showing majority U->C changes (excess C->U) were expressed as proportion of the number of variable sites in the alignment, with totals sub-divided into sites showing different ranges of AI values (Fig. 9). There was a substantial over-representation of sites showing excess C->U mutations in HCV, HPgV and other virus datasets showing the C->U / U->C asymmetry, particularly at sites with high AI values. From these, it appears that the asymmetric mutational process contributes a substantial proportion of their standing viral diversity in all four of the viruses analysed (≈ 15% of variable sites).

**FIGURE 9.**
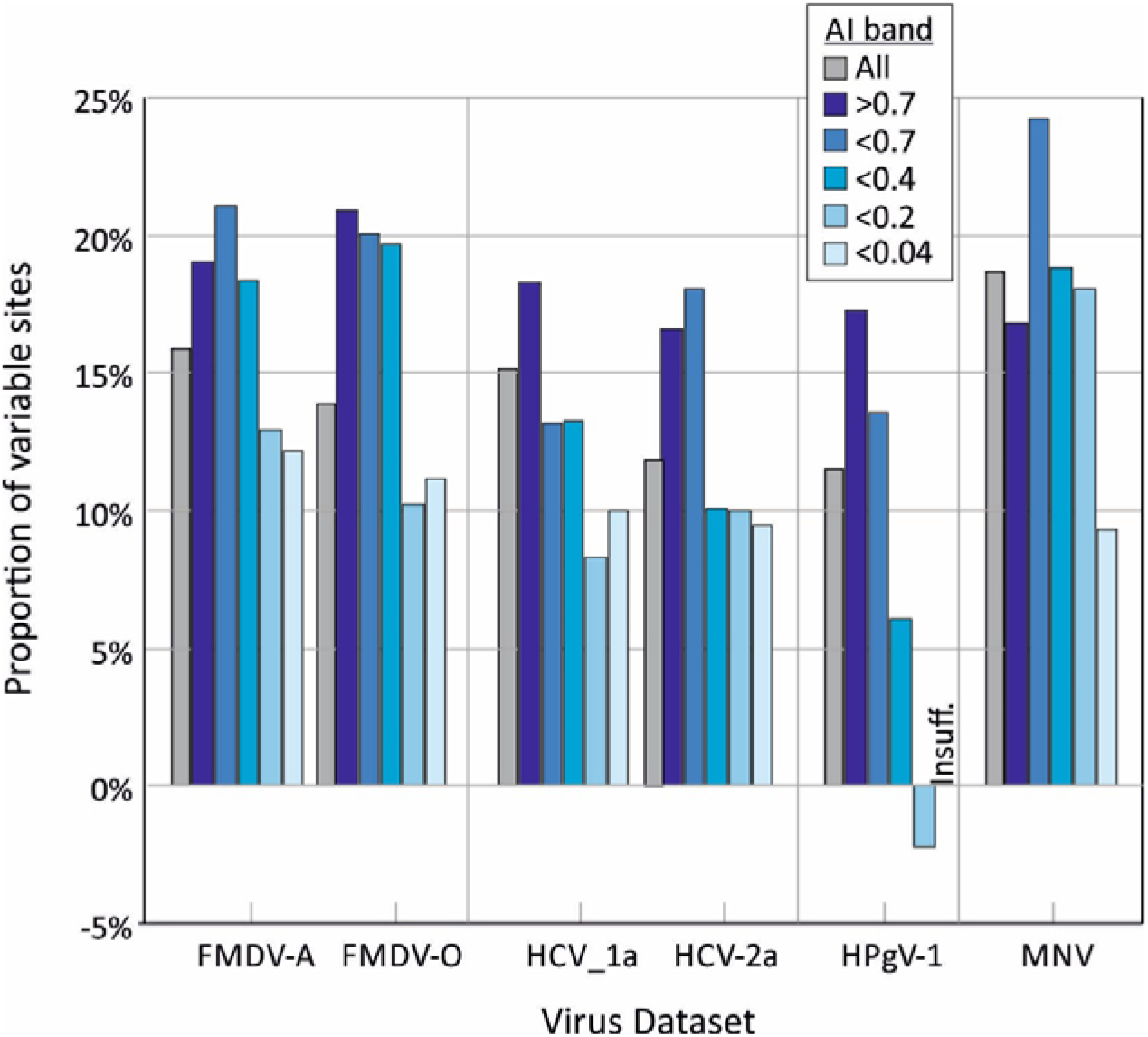
PROPORTIONS OF SITES MUTATED BY C->U / U->C TRANSITION ASYMMETRY. Excess C->U mutations (number of sites with majority C->U transition – sites with U->C) expressed as a proportion of all variable sites in genome alignments of viruses showing C->/U->C asymmetry. Variable sites were further sub-divided by AI band.

## DISCUSSION

### C->U transition asymmetries

This study provides evidence for an excess of C->U changes over the reverse (U->C) and complementary (G->A) transitions in a diverse range of RNA viruses, including HCV (all genotypes), FMDV, MNV, HPgV-1 and rubella virus. Fold excesses ranging from 1.9x – 3.6x (Table 1; Fig. 3A) approached those reported in previous analyses of SARS-CoV-2 (7.5x), SARS-CoV (4.2x), MERS-CoV (3.0x) and were comparable to those reported in seasonal coronaviruses (1.8x – 3.0x) (21). There has been little investigation or analysis of the phenomenon of mutational asymmetry in RNA viruses, although a recent study identified a noticeable excess of C->U changes in the genomes of RUBV persisting in patients after immunisation (24, 31). Using the analytical methods of the current study on that dataset, RUBV sequences showed a C->U / U->C transition asymmetry (base-corrected) ratio of 1.7 (data not shown), while the analysis of a larger dataset of circulating rubella virus strains (n = 73) in the current study produced an even higher C-> U / U->C transition asymmetry (3.6x; Fig. 4). Although the authors linked the asymmetry to RUBV persistence (24, 31), the finding of even greater asymmetry in circulating RUBV strains indicates this phenomenon occurs as part of a natural transmission chain of rubella virus infection.

### Mutational mechanisms

The directionality of the observed C->U transitions provided evidence for a strand-specific mutational process. As discussed previously (21), this would not occur as a result of systematic transcription errors by the viral RNA dependent RNA polymerase during virus replication. Progeny viral RNA genomes are generated, by definition from an equal number of positive and negative strand copyings, and a tendency to misincorporate a U instead of a C would be reflected in a parallel number of G->A mutations where it occurred on the minus strand. However, as observed previously for SARS-CoV-2 (21-24), the frequency of G->A mutations was substantially lower than C->U changes, and generally comparable to those of the other transitions, A->G and U->C (Figs. 1, 2).

It could be argued that natural selection operates differently on the minus- and plus-RNA strands where the limited number of minus-strand templates produces a much larger number of genomic RNA. There may be more stringent selection against mutations occurring in the negative-strand as these are invariably copied into the plus-strand, whereas plus-strands are relatively dispensable and mutations occurring during their synthesis will not overly affect the overall replication process. Indeed, full-length transcripts with plus-strand synthesis errors could be readily packaged into virions assembled from viral proteins synthesised from independently transcribed mRNAs. However, for there to be a strand-specific imbalance in the number of C->U mutations, there would have to a particularly elevated frequency of C->U transcription errors associated with the SARS-CoV-2 RNA dependent RNA polymerase (RdRp). Enzymatic data on error frequencies in viral RNA polymerases is relatively scarce. However, biochemical assays of misincorporation kinetics on RNA templates by the poliovirus RdRp (32, 33) or retroviral reverse transcriptase (34) did not identify C->U misincorporation to be favoured over other mutations although it is conceivable that other RNA virus RdRps may possess distinct mutational spectra.

The validity of this alternative account of the strand-specificity of C->U transitions depends also upon the testable prediction that natural selection on the minus-strand would favour fixation of mutations that were phenotypically neutral over non-synonymous mutations. C->U strand asymmetry should therefore be reduced at synonymous sites. In practice, however, analyses of SARS-CoV-2 genomic variability shows that synonymous C->U transitions dominate the SARS-CoV-2 mutational spectrum when these are analysed separately from non-synonymous substitutions (35) and are over-represented in recurrent homoplastic mutations (36).

<Why would there be a 5’ context of based on differential selection?>

Editing of viral RNA sequences by ADAR1 has been documented to occur extensively during the replication of a number of RNA viruses (37), but as its substrate is double-stranded RNA, similar editing activity on both strands would not create mutational asymmetry. Furthermore, its mutation of A to inosine which is then read as a G is the opposite of what is observed in the RNA virus datasets analysed in the current and previous studies (21, 23, 24). The tentative explanation for the observed C->U editing in coronaviruses is that it is driven by one or more of the APOBEC proteins that catalyse C->U mutation on a single stranded template (21, 23, 24). However, this hypothesis requires functional investigation; It should, for example, be reconciled with the previous finding that antiviral effects of APOBEC occurred in the absence of any evidence for RNA editing of coronavirus HCoV-NL63 genomic sequences (38). In contrast to the abundance of evidence for APOBEC3-induced mutations as a cellular defence against retroviruses and DNA viruses, further studies are required to determine whether RNA viruses can be functionally restricted by similar mechanisms.

Inferences on mechanism based on bioinformatics analyses therefore must at this stage be indirect; they should additionally acknowledge that the currently recognised repertoire of RNA editing pathways and other mutational mechanisms in mammalian and other vertebrate cells is almost certainly incomplete. Analysis is further complicated by the diversity of the APOBEC gene locus, often quite different editing substrates and favoured contexts between paralogues and substantial differences in APOBEC gene numbers between mammalian species. In terms of which paralogs may mediate RNA virus editing, several authors have proposed APOBEC3A (A3A), based on its previously demonstrated ability to mutate cellular mRNAs and a context preference for a 5’ U upstream of the editing site (39, 40). Alternatively, the cytoplasmic location, induction by interferon and evidence for positive selection and diversification during mammalian evolution, A3F and A3G (38, 41, 42) or APOBEC1 in a rat (43) have been proposed as additional candidates, although the 5’ context preference of A3G for a 5’C conflicts with previously documented higher C->U transition rates of edited RNA in 5’ U or 5’ A contexts (44, 45). The increased C->U/U->C mutational asymmetry downstream of a U observed in all RNA viruses after normalisation for differences in G+C content (Fig. 5) is consistent with the RNA editing preferences of human cellular mRNAs by A3A (40, 46) and in a previous analysis of rubella (31).

Where and how editing of RNA virus genomes might occur in the infected cell is unclear. Strand-specificity and its presumed requirement for ssRNA sequences suggests that editing takes place on naked genomic RNA, perhaps during virus entry or during packaging. As APOBECs are IFN-induced, editing during packaging appears more likely notwithstanding the fact that coronavirus RNA genomes, like –strand RNA viruses, are typically associated with ribonucleoprotein throughout the replication cycle and must substantially limit their exposure to host pattern recognition receptors. Finally, it need not be assumed that C->U editing in virus genomes may occur through the action of only one APOBEC protein or indeed may originate from an entirely different mechanism outside the documented RNA editing functions of ADAR1 and APOBECs. As suggested by the authors, overlapping but nevertheless distinct C->U mutational contexts in SARS-CoV-2 and rubella genomes points towards the operation of more than one mutational mechanism (24); the heterogeneity in the effects of 3’ base contexts (Fig. 5) may be further evidence of the operation of different mutational mechanisms. Establishment of effective methods to induce and quantify RNA virus editing *in vitro* and a targeted gene deletion approach to functionally test editing abilities of individual APOBEC proteins on RNA templates would be important steps in such investigations.

### Influence of RNA structure on editing

Apart from 3’ and 5’ base context, the extent of C->U / U->C provided evidence that a substantial proportiontransition asymmetry was not influenced by any standard RNA composition metrics, including G+C content and CpG or UpA dinucleotide suppression (Table 2). Unexpectedly, however, the phenomenon appeared to be largely restricted to RNA viruses with the previously described property of genome-scale ordered RNA structure (25, 26, 47), that refers to the presence of pervasive RNA folding throughout the genomes of a subset of vertebrate RNA viruses. There was a significant association between MFED value and C->U / U->C asymmetry ratio (Table 2; Fig. 3B), but no association with the G->A / A->G normalised asymmetry values.

Intriguingly, the observed association of C->U mutational change appears to be paralleled by the previous noted restriction of editing of human mRNA sequences by A3A to sites possessing defined RNA secondary structure (46). In an analysis of human transcriptome RNA sequences, over half of the identified edited sites were flanked by short palindromic sequences, typically locating the edited base in the terminal unpaired region of a stem-loop. Supporting this RNA structure association, an analysis of the contexts of SARS-CoV-2 and rubella C->U edited sites identified preferential C->U mutations in terminal loop compared to stem sequences (24), or in predicted unpaired compared to unpaired regions (27). This association may explain why coronaviruses, with their consistently high MFED values and intensely folded genomes (27, 48, 49) may be particularly targeted by RNA editing. The observed association of excess C->U changes in HCV and other structured viruses is consistent with the action of one or more APOBEC isoforms directly editing a wider range of RNA virus genomes.

### Evolutionary consequences of widespread RNA editing

The analyses performed in the study provided evidence that a substantial proportion of population variability in HCV and other structured viruses could be attributed to a C->U hypermutation phenomenon (15% - 20% of total sequence changes; Fig. 9). The changes correlated poorly with overall phylogeny of viruses in the alignment based on association index calculations (Fig. 7A), consistent with previous analyses that documented extensive homoplasy of C->U changes in SARS-CoV-2 (21, 22). The limitations of this type of analysis should however be acknowledged. While the association index calculation represent an established and robust, phylogeny-based method for evaluating group membership with phylogeny (50, 51) and performs well compared to other metrics of genetic partitioning (52), analyses in the current study were based upon groupings derived from base identities. These are necessarily limited to between two and four character states defining groups at each alignment position, and this restriction may create mutational saturation effects at highly variable sites and lead to false detection of homoplasy. However, even sites with relatively low Shannon entropy values (0.3-0.5), that would not create any consistent saturation effect showed an excess of C->U changes in HCV and HPgV-1 alignments (Fig. 8).

The distribution of C->U changes (and other transitions) at an individual sequence level approximated to a Poisson distribution (Fig. 3), albeit with some over-dispersion in the EV-A71 sequence dataset arising from its phylogenetic structuring (Fig. S2; Suppl. Data) compared to HCV.There was no evidence for the occurrence of individual hypermutated sequences in a background population of non-mutated sequences, as previously observed in HIV-1 and hepatitis B virus (HBV) (17, 28, 29, 53, 54). This likely originates from the differences in replication cycles of RNA viruses and retro-transposing viruses – the RNA virus sequences in the current study were derived from consensus sequences and therefore would have to be fully replication competent and evolutionarily fit to be represented in clinical samples. In contrast, hypermutated sequences of HIV-1 are typically derived from integrated proviruses derived from APOBEC-edited reverse transcription. Their survival in memory T cells occurs irrespective of whether they are able to generate infectious virus or not; similarly for HBV (53, 54). Such sequences can therefore accumulate extensive and bizarre mutational damage as they are effectively evolutionary dead-ends. In marked contrast, the excess of C->U changes edited into RNA virus genomes susceptible to editing may therefore represent the maximum tolerable mutational load compatible with viability and onward transmission.

While tangential to the primary focus of the study, the shape of phylogenetic trees constructed from different RNA viruses differed substantially (Fig. 8; Fig. S1, Suppl. Data) and potentially contributed to observations of homoplasy. These differences in branching density have been previously quantified using the temporal clustering (TC) metric (55). The bush-like, over-dispersed topology of HCV showed a lower TC value that derived from a neutral evolutionary simulation, a difference attributed to potential rate variation in different lineages of HCV or population subdivision which promotes the co-existence of lineages. The latter model may potentially be equated with distinct patterns of endemic and epidemic partitioning in the different trees associated respectively with persistent and non-persistent virus infections (Fig. S1; Suppl. Data). However, there is the further possibility that tree shape and the associated occurrence of phylogenetically uninformative sites in structured virus genomes may also be influenced by extensive RNA editing and homoplastic cycles of mutation and reversion as observed in SARS-CoV-2 (21, 22). The development of evolutionary simulation methods where RNA editing is incorporated and parameterised may lead to valuable insights into the nature and trajectory of short-term diversification. It may serve to better characterise the evident differences between RNA viruses in the nature of their divergent evolution. The observation that excess C->U changes accounted for 15%-20% of variable sites of HCV, HPgV-1, FMDV and MNV at any one time (Fig. 9) indicates the powerful role that RNA editing may play in the generation of RNA virus diversity.

## MATERIALS AND METHODS

### Sequence datasets

Alignments of sequences of HCV genotypes 1a, 1b, 2a and 3a, SARS-CoV-2 and other coronaviruses were derived from previous studies (27, 47). Further alignments of other RNA viruses were constructed for the study from GenBank and the VIPR database (56) using all available or randomly selected sequence subsets as described in Table S1; Suppl. Data. Coding region sequences were aligned using MUSCLE (57) as implemented in the SSE package version 1.4 (http://www.virus-evolution.org/Downloads/Software/) (58). Analysis of viruses encoding single polyproteins (*ie*. picornaviruses, flaviviruses) was based on coding regions only. Regions spanning the start of the first open reading frame (ORF) to the end of the last ORF were used for analysis of viruses with polycistronic genes (coronaviruses, togaviruses, pneumoviruses, filoviruses, hepeviruses and caliciviruses). Alignments are available from the author on request.

### Sequence analysis

Calculation of pairwise distances and nucleotide composition was performed using the SSE package version 1.4. Sequence changes were compiled using the program Sequence Changes with a variability threshold typically set at 5% heterogeneity, where heterogeneity was calculated as the cumulative frequency of all non-consensus bases. Multiple thresholds were used to analyse mutation representation at sites showing different levels of variability (Figs. 1, 2). RNA secondary structure predictions used for computation of MFE and MFED values was carried out using the program Folding energy scan in the SSE package using sequential 300 base sequence fragments incrementing by 30 bases between fragments. The program call the RNAFold.exe program in the RNAFold package, version 2.4.2 (59) with default parameters.

### Association index calculations

AI values were calculated using the algorithm originally described by Wang *et al*.(50) and Cochrane *et al*. (51) and implemented in the SSE package. The assignment of group labels based on nucleotide identity at sequential sites in an alignment was automated in the program extension, homoplasy scan in the SSE package version 1.5.

### Phylogenetic analysis

Neighbour joining trees were constructed from aligned sequences using the program MEGA7 (60). Lineage against time plot were derived from data generated in the Phylocom package (30).

### Statistical analysis

All statistical calculations and histogram constructions used SPSS version 26.

## ACKNOWLEDGEMENTS

The work was supported by a Wellcome Investigator Award Grant WT103767MA to PS. MAA is supported by a Wellcome Sir Henry Dale Fellowship (220171/Z/20/Z). We are very grateful to Jeremy Ratcliff for critical reading of the manuscript.

## SUPPORTING INFORMATION

Table S1. Accession numbers of sequences in virus alignments

Table S2. Compositional features of RNA virus sequence datasets used in the study

Figure S1. Unrooted phylogenies of sequence alignments used for homoplasy analysis

Figure S2. Lineage through time plot for sequence alignments used for homoplasy analysis

