## Supplementary Data for "Mutation bias implicates RNA editing in a wide range of mammalian RNA viruses"

TABLE S1

ACCESSION NUMBERS OF SEQUENCES IN VIRUS ALIGNMENTS

**RSV, genotype A	VIPR Database	Selection of 100**

KM517572, KJ672442, KJ627291, KU316131, KP258740, LC474558, MG642063, KU950502, KX765898, KU950538, KJ627690, MG642032, MN630107, KX655627, KU950474, KU950671, KX765887, KX655692, MH447956, MK167036, KU316137, KU950537, KJ723489, KU950583, MG793382, KU950520, MF001043, MF614946, MG642033, MN630088, KJ672471, MK749909, KP663728, MG642052, KX765934, KU316162, KJ672479, KJ672427, KU950642, KM578843, KJ939957, JX069802, KP258700, JQ901450, KJ672449, KJ627687, KU950652, KJ723474, KX655689, KU950685, KM042390, KJ627273, KJ627287, MG642038, MN306030, KU316168, KX655652, KY883567, KJ939936, KJ672482, KF973330, KU316164, KJ672455, KP258695, KJ939964, KY982516, KJ627733, KX655626, KP258696, KP258704, KP317953, KU950594, MN306054, KX765904, KX655670, MK109776, KJ643531, KX765942, KU950686, KU950501, KJ643465, KJ627343, KU950634, KP258733, KJ627258, KJ627674, MN630102, KJ627719, KF973329, KJ643503, KJ723473, KP258701, KU950541, KJ627723, KU950590, KJ627372, KJ627708, KU950595, KJ627325, KP218910

**BVDV	VIPR Database	All (no insertions)**

KC853441, MH490942, HV302650, DD122170, AX741975, AF091605, AJ585412, M96751, AJ133739, KY964311, AB078950, MH133206, MH231153, JN704144, KC963967, LC068605, LC068604, LR760748, MH899944, LR699800, LT837585, KP941584, AF220247, LR699799, KR866116, MT079816, MN188073, KF501393, KC695814, KT355592, KC853440, JN400273, MK102095, MH490943, KT896495, LC089875, U86600, KC757383, LC089876, KF772785, KJ620017, KP313732, AF041040, KX987157, LT907992, KX577637, MF166858, MG923683, KP941589, MH899943, EF101530, U63479, KP941588, KY849592, KX857724, DQ088995, KP941581, LT968777, KP941591, KP941590, KP941583, MN188074, KT951841, KT951840, AF526381, KU159365, KT943518, KJ689448, KR029825, JQ799141, KP941592, LT631725, MH899941, MT265675, KF896608, MH166806, KP941587, MH379638, KJ541471, MH899942, MF693403, MK509773, MH899945, KP941586, LT797813, MK509775, KC695810, MK509774, MN623291, KF835699, JX419398, JX419397, KF835697, KF835698, LR699802, LR699801, KU756226, LR699803, MF278651, JQ418633, HQ174296, HQ174295, JN380083, HQ174294, JN644055, HQ174293, JN380080, HQ174292, MF278652, JQ418634, MF172980, JN380089, JN380088, JX297517, JX297521, JX297520, JX297519, JX297518, JX297516, JX297515, JX297514, JX297513, JX297512

**HPeV, genotype 3	GenBank	All**

KX068679, KM986843, KJ659490, AB084913, AJ889918, KY020128, JX682576, GQ183029, GQ183033, GQ183032, GQ183030, GQ183031, GQ183026, GQ183027, GQ183028, KY556667, KY556665, KY556659, KY556669, KY556666, KY556664, KY556660, KY556672, KY556662, KY556670, KY556661, KY556668, KY556663, KY556674, KY556675, KY556671, KY556673, MK604044, MK604037, MK604067, MK604040, MK604047, MK604042, MK604039, MK604061, MK604059, MK604062, MF371334, MF371336, MF371333, MF371335, MK604049, MK604046, MK604048, MK604060, KT879915, KT879922, MK604043, KY351616, KY351631, KY351617, KY351618, JX826607, MK604058, MK604052, MK604057, MK604041, MK604045, MK604063, MK604066, MK604038, MK604051, MF797924, MF797928, MF797929, MF797934, KT626009, MF797935, MF797930, KU556748, KU556749, MF797933, MF797926, KY351632, MF797927, MF797925, KU556750, MF797932, MF797916, MF797931, AB668029, AB668030, AB668032, AB759186, AB759187, AB668033, AB668031, LC043117, LC043115, LC043116, LC043114, LC043128, AB759185, AB759191, LC043121, AB759189, LC043127, LC043124, AB759190, LC129275, LC129269, LC129276, LC129272, LC129271, LC129270, LC129273, LC129274, LC383404, LC383392, LC383394, LC383405, LC383407, LC383409, LC383399, LC383408, LC383393, LC383396, LC383400, LC383402, LC383410, LC383391, LC383406, LC383403, LC383401, LC467800, AB759205, AB759204, AB759206, AB759200, AB759201, AB759202, AB759203, AB759199, AB759197, AB759207, AB759192, AB759198, AB759193, AB759195, AB759194

**CHIKV	GenBank	All**

HM045811, HM045792, HM045810, KX262990, HM045809, HM045823, HM045821, HM045813, HM045786, HM045816, HM045785, HM045788, KY038946, HM045814, HM045805, HM045822, HM045808, HM045815, HM045818, HM045812, KY038947, KX262995, KX262986, HM045791, HM045784, HM045797, HM045800, HM045790, HM045806, KX262988, HM045789, HM045819, HM045820, HM045796, HM045787, KX262987, KX262991, KP702297, KF283986, FR717336, HM045817, JF274082, EF210157, KP003808, KJ941050, HM045794, FJ807896, GQ428211, FJ959103, FJ000068, FJ000067, FJ000066, FJ000065, FJ000064, FJ000063, FJ000062, EU564335, EU564334, KP003809, KP003807, GU189061, GQ428210, KX262996, EU703761, FN295483, FN295484, EU703760, EU703759, GQ428213, GQ428212, KX262993, KX262989, KP003811, KP003810, HM045799, FJ445428, FJ445427, FJ000069, EU244823, EU372006, HM045801, KP003812, KM923918, KM923917, FJ807897, GU199351, FR687343, FR687342, FR687341, FR687340, KJ796851, KJ796849, KJ796848, FN295485, FN295487, JN558835, GU301780, GU013528, GU013529, GU013530, FJ445502, FJ445484, FJ445463, FJ445432, FJ445430, FJ445426, GU199353, GU199352, FJ807898, GQ428215, GQ428214, FJ513675, FJ513673, FJ513657, FJ513654, FJ513645, FJ513637, FJ513635, FJ513632, FJ513629, FJ513628, FJ513679, FJ807899, FJ445511, GU199350, KJ796844, KF151175, KX262997, FR687348, FR687347, FR687346, FR687345, KJ796852, KJ796850, KJ796847, KJ796846, KJ796845, KF151174, JN558836, GU301779, GQ905863, GU908223, KP164869, FR687344, GU301781, KT336777, JN558834, HQ846358, JX088705, HQ846356, KF590567, KF590566, KF590565, KF590564, KC862329, HQ846359, KT336778, JQ861256, JQ861260, JQ861259, JQ861258, JQ861257, KJ679578, KC614648, KJ679577, KP003813, HE806461, KU365292, KT336780, KT336779, KF318729, KC488650, KT308163, KT308161, KT308162, KT308160, KT308159, KT336782, KT336781, KJ579187, KJ579186, KJ579185, KJ579184, KJ689453, KJ689452, AB860301, KJ451623, KJ451622, KM673291, KF872195, KP164570, KP164569, KP164568, KR559482, KR559486, KR559493, KX702402, KR559476, KX702401, KX262994, KX262992, KT192707, LN898111, LN898110, LN898108, LN898107, LN898103, LN898101, LN898100, LN898098, LN898095, KR046234, KR046233, KR046232, KR046230, KR046229, KR046228, KR046227, KR559498, KR559496, KR559495, KR559494, KR559492, KR559490, KR559488, KR559487, KR559485, KR559484, KR559483, KR559481, KR559475, KR559474, KR559470, KR264951, KR264949, KP164571, KP164567, KP851710, KP851709, KR264950, KU940225, KR559473, KY057363, KX228391, KX496989

**EBOV	GenBank	Selected**

AF086833, KC242801, KY425630, KY425637, KY425639, KY425647, KY425649, KY425652, KY425656, MH121166, MH121168, KR063671, KC242791, KC242792, KR063672, JQ352763, KR867676, KU182898, KU182899, KU182900, KU182901, KU182902, KU182903, KU182904, KU182905, KU182906, KU182907, KU182908, KU182909, KY425636, KY425653, MG572235, KC242796, KC242799, KR824526, MH121165, KC242793, KC242794, KY785939, KY785940, KY785947, KY785948, KY785949, KY785965, KY785969, KY785970, KY786017, KC242800, KF113528, KC242784, KC242785, KC242786, KC242787, KC242788, KC242789, KC242790, HQ613403, MH733477, MH733478, MH898466, MH733479, MH733480, MH733481, MH733482, MH733483, MH733484, MH733485, MH733486, MH733487, MH733488, MH733489, MH733490, MH733491, MK007329, MK007330, MK163644, MK163645, MK163646, MK163647, MK163648, MK163649, MK163650, MK007331, MK007332, MK007333, MK007334, MK007335, MK007336, MK163651, MK163652, MK163653, MK163654, MK163655, MK163656, MK163658, MK163660, MK731985, MK731987, MK731988, MK731993, MK007337, MK007338, MK007339, MK007340, MK007341, MK007342, MK007343, MK007344, MK731986, MK731989, MK731990, MK731994, MK163661, MK731992, MK163662, MK163663, MK163665, MK163666, MK163667, MK163668, MK163669, MK163670, MK163672, MK163673, MK163674, MK163675, KU143777, KT725268, KX009901, MF599519, KT725257, KR105295, KR653240, KR534585, KP759733, KP759660, KT725311, KR817226, KR817112, KM233085, KR534564, MH425138, KY426709, KR075003, KR105300, KR817224, KP759692, KY401669, KP184503, KR534517, KT357815, KR817200, KU143821, KY558986, KU143778, KT725380, KX121424, KR653304, KR817126, KM233088, KU143812, KR105249, KY471125, KR817222, KM233106, KM233058, KX009897, LC152433, AY354458, KP759674, KU052670, KR534528, KP759724, KM233116, KR653279, KT357841, KP759728, KR105323, KM233050, KR653301, KY007525, KT725296, KR817082, KU143833, KR105316, KR105244, KR534574, KR105283, KY426704, KY471092, KR534546, KP728283, KR534552, KT725349, KU296527, KM233087, KT013257, KR817111, KT725319, KR105232

**MeV	GenBank	All**

AF266288, AY730614, JF727649, AY486083, AF266290, AF266289, AB591381, AF266287, AF266286, Z66517, EF033071, AB046218, FJ416067, S58435, MN145872, K01711, EU435017, DQ345721, MG912589, AB016162, AB032167, DQ227321, JF791787, LC336599, EU293548, EU293552, EU293551, DQ227319, EU293549, AB481087, AB012948, AB481088, AB012949, DQ227318, JN635404, JN635407, DQ227320, MN131123, MH356254, JN635402, MH356236, KY969480, MN125020, HM439386, MG912590, JN635403, JN635410, MH356241, MH356250, MF496202, MF449469, KC164757, MN125028, MH356253, MN131117, MF496200, MH356244, MH631015, MH356239, MF496201, MH631016, MH356243, MG912593, MH356242, GQ376026, MK513625, MH356240, MH638233, GQ376027, MH173047, JN635405, MH356245, JN635406, KJ410048, MK161348, MG912592, JN635408, KY969479, MH356248, KJ018971, MH356237, AB254456, KU728742, JN635409, KU728743, MT738550, MH356252, MK142916, MK142914, MK142915, MH356238, MH356246, KT732231, MH356247, KC117298, MH356255, MH356249, MT789844, MT789833, MT789849, MT789848, MT789839, MT789838, MH356251, MT789842, MT789840, MT789832, MT789824, MT789827, MT789821, MT789847, MT789829, MT789825, MT789823, MT789836, MT789830, MT789837, MT789826, KY969477, MT789828, MT789835, MT789843, MT789834, MT789822, MT789831, KT732230, KY656518, KT732225, MT789845, KT732258, KT732254, MT789841, KT732244, KT732237, KT732234, KT732233, KT732227, KT732226, KT732256, KT732246, KT732240, MT789846, KT732235, KT732228, KT732241, KT732251, KT732232, KT732229, KT732249, KT732247, KT732243, KT732252, KT732255, KT732245, KT732260, KT732259, KY969476, KT732261, KX838946, MN630022, KY969478, MG912591, KT732214, KY969481, MN017369, MN893225, KC164758, MN630023, KT732219, KT732224, KT732222, KT732221, KT732217, KT732223, KT732216, KT732215, MF775733, KT732220, KT732218, MT789797, MT789815, MT789818, MT789805, MT789804, MT789800, MT789799, MT789795, MT789793, MT789792, MT789813, MT789796, MT789814, MT789803, MT789802, MT789801, MT789798, MT789790, MT789789, MT789788, MT789807, MT789806, MT789816, MT789812, MT789811, MT789791, MT789808, MT789817, MT789810, MT789819, MT789820, MT789794, MT789809, MK513610, MK513616, MG912594, KJ755975, FJ161211, DQ211902, JN635411, KJ018970, MN630021, MN630020, MG972194, KT588921

**EV-A71	GenBank	All**

MK652139, DQ341367, MG976581, KX372324, JX244184, MH716347, JQ639384, LT719065, LC506514, KC436271, MG214681, KT428646, KP289432, MH716380, JF738002, HQ647173, GQ279369, KU936120, HM807310, FJ607337, FJ357384, MG367595, LC321989, MH716337, AB575917, KX372322, KJ004560, KF982854, JQ074189, MG367594, EU131776, LT719063, LT719067, AB575918, MN254979, MF662695, KJ400360, U22522, KJ686246, KU936132, AB575938, HM002487, HQ647172, KJ686249, JN992284, KC436265, AB575935, KU936128, FJ194964, KJ784495, AB747375, JX678885, DQ341358, LC375764, AB575928, MF973167, LC506513, FJ357376, LK985324, MG672479, KF142413, MG773126, KX372317, KP289419, KP289425, KP289422, AB575911, KT345959, MG672481, GQ994992, DQ341364, DQ341357, KU936121, KC436266, KY612315, KC109780, AB575914, GQ231939, MH111073, JQ681218, KJ686308, MH716310, AF302996, GQ994989, KY315729, AM396584, LT719064, KM055005, KU254596, KP308450, HQ456306, KC436270, MG367600, KU936125, JQ742001, KC954664, FJ607336, HQ189392, KT008669, DQ341355, JQ742002, KX372328, JQ280307, KX372310, KM673244, DQ341359, KT428647, AM396587, KJ686262, FJ606447, KP289431, KX197462, AB575936, MG672478, LT719068, KP289424, EF063152, MG207962, EU414333, FJ360545, AF316321, JF738001, MF662701, KJ686208, KP308426, JF799986, MH484070, KF974790, GQ231934, U22521, AB575923, GQ994991, MG431943, JQ316638, DQ341360, KX372308, AY465356, HQ647179, LC506515, LC506516, AB204852, KF501389, AB204853, GU434678, MH484071, MH484069, AB482183, MH484068, AB747373, MH484067, MH484066, KX139462, MG432108, KF514878, AJ586873, GQ231936, FJ357385, GQ231942, KT354870, KT354867, KF974789, LC321990, GQ231935, KX372330, KT354869, KT354868, KT354866, KF974784, MH716379, KF974779, GQ231943, GQ231925, MG756712, KT354872, KT354871, MG756729, KU641504, KU641502, GQ231941, KT354875, KT354874, MH716377, MG756708, KX372314, KU641505, KX372327, KF154355, MG756743, MG756709, AB575913, MG756713, LC375765, KF974794, AB550337, MG756714, KU641503, KF974781, KF974797, KF974792, MG756749, KU641507, KU641501, MH716376, MH716386, KU641506, KF974780, MG756710, MH716384, KX372316, MH716348, KX372323, AB575927, MH716391, MG756734, KF974785, KF974783, MH716389, MG756740, FJ357375, MH716383, MG756750, MG367599, MG756742, MG756733, MG756727, MG756723, MG756721, KX372331, MG756752, HQ647171, MG756746, KF974787, MG756741, MH716390, KU641508, MH716353, MH716351, MH716300, MH716299, MG756744, MG756730, MG756737, KF974796, KF974791, MH716361, MG756747, MH716350, MG367598, AM396585, MG756753, KY074644, MH716362, MH716349, KP691653, MH716360, MG756736, MG756735, MG756745, FJ357382, MH716319, MG756726, MH716364, KP308454, KF974793, MH716292, MG756751, MG756706, FJ357380, KJ686176, KF974795, MH716385, KP691659, KJ686137, MH716366, MG756728, JX678878, MG756711, AB550333, MG756732, JX025561, MH716288, MH716286, MG756748, KX372326, JX678886, DQ341362, MH716388, KX430824, MH716346, MG756738, KM508794, KJ686302, KJ686131, MH716387, KP691660, GQ231930, GQ231928, EU527985, MN629889, MH716287, MH716332, MH716313, MH716304, KJ686225, MH716393, MH716311, AB550338, FJ607335, MH716352, KJ686270, GQ231929, GQ231927, MH716267, EU753365, MH716305, MH716285, AB550339, MH716284, JX678882, KJ686128, GQ279370, MH716298, MH716259, MH716290, MH716282, MH716281, KP691643, MH716344, JQ965759, KF974798, KF154354, MH716315, EU753407, MH716291, MH716272, MH716271, MH716373, MH716274, KJ686211, KJ686264, AF176044, MH716318, MH716317, MH716314, KP691657, MH716334, MH716270, MH716293, MH716278, KP691663, JQ517316, FJ357374, KF514880, MG756731, KX372311, JN544418, EU753375, HM622390, AF304458, MH716283, KP691648, MH716381, MH716333, MH716265, FJ713137, MH716260, MH716356, MH716268, MH716266, AF352027, MH716280, KJ686222, MH716308, MH716369, KY952186, KR045298, EU753398, MH716269, KX372320, KP691654, JQ708210, JN020147, MH716273, HQ611148, MG756739, KJ686140, AF136379, MH716277, KP691647, MH716302, MH716297, KP691658, MH716374, MH716336, MH716258, KJ686192, MH716316, KX372313, KP691664, JN992282, JN544419, MH716303, KP691652, GQ994988, KJ686297, MH716261, MH716309, GQ231926, MH716279, KP691651, MH716331, KP691650, AB550341, MH716342, GQ231940, MK904809, KP691655, AF304459, MH716324, MH716294, GQ231937, GQ231932, DQ341365, JQ724182, JQ074188, KF134486, MH716275, GQ231931, MH716358, MG756704, JN864018, MH716323, MH716276, KP691662, MH716355, MH716327, MG756698, KU936130, KX197459, JQ074187, KP308430, JN992285, MF662687, MH716296, JN256063, KP308419, KJ686234, AB469182, JX678877, JX678876, AB550334, EU703813, JN256064, AB550335, KM077140, EU376004, MH716325, KJ004556, HM053669, MH716328, KJ746494, KX197457, JX678883, DQ341368, MG756696, EU414334, JX986739, HQ891924, FJ606448, EU703814, KX197463, KP308449, MH716329, MH716252, EU376005, EU364841, MG756702, KY074643, KP691645, JF830007, GU396280, FJ158601, MH716335, KC436269, HQ891923, EU864507, MH716257, KC954663, DQ060149, FJ606450, EU414335, EU703812, DQ381846, MH716354, KP691665, KY425527, KJ686195, HQ456311, FJ194965, MH716321, MH716256, MH716341, KP691656, FJ158600, MH716370, JX025559, FJ607338, KR045302, KP308459, KP308428, KJ686226, KU574619, KR045301, JX678884, JX678880, JX678875, MF662686, HQ456305, AM396586, JN256060, HM002489, KP308434, JX111890, AM396588, JQ708209, JX111892, HM002485, AF119796, MG756725, KX372318, JX111888, MH716248, MG756697, MG756700, JX678881, HQ694982, MG756699, KJ686237, KJ686198, EU414331, MF662685, MH716254, KJ686269, KJ686200, KJ686194, KJ686157, MF662681, KY014080, KJ686184, JN256059, HQ891927, JX244183, KP308457, KP308401, GU459071, KP308409, KJ686307, KJ686261, KX752783, KP308453, KP308441, KJ686204, KJ686151, HM622392, MH716255, KP308408, KJ686280, KJ686256, KJ686248, KJ686166, HQ456310, KJ686224, KJ686223, JF913464, KX372332, KX197458, KP308442, KP308436, KJ686238, MF662684, MH716340, MH716253, KP308439, KJ686206, JX111891, MH716251, KP308407, JX244186, FJ439769, MG756722, HQ694985, KP308437, KJ686245, KJ686168, KJ686142, KJ686127, JX244185, HQ891928, KP308432, KJ686266, KJ686217, KJ686201, KJ686165, KJ686295, FJ360546, KP308415, GU459070, KP308460, KP308440, KJ686258, KJ686182, KJ686162, KJ686143, JX111893, JX017384, KT008671, KP308412, KJ686299, MH716250, KP308422, KP308402, KJ686284, KJ686252, KJ686251, KJ686163, KJ686152, HQ891925, FJ600325, KJ686268, KJ686185, KF826491, KT008672, KJ686193, KJ686305, KJ686229, KJ686239, KJ686236, KJ686170, KJ686156, KJ686149, KJ686141, KJ686279, KJ686216, HQ694983, KP308452, KP308438, KJ686241, HM622391, KP308403, KJ686303, KJ686283, KJ686267, KJ686263, KJ686231, KJ686213, KJ686210, KJ686155, KJ686132, KJ686130, KJ686221, KJ686188, KJ686287, KJ686174, KJ004559, KJ686286, HQ647177, KP308433, KJ686214, KJ686202, KJ686161, KJ686160, KR045300, KJ686175, EU414332, HQ882182, HQ694986, HQ694984, HM003207, MK028135, HQ647178, KJ686196, KJ686172, MF662680, KJ686289, MG773125, KY014079, KT008670, KJ686197, KJ686153, KP308423, KJ686259, KJ686171, KP308451, GU198368, KT428649, KJ686300, MF662678, KP274877, KJ686298, KJ686181, KJ686158, GU198369, KJ686189, KJ686129, GU198367, JF894383, JF894381, KJ186973, KJ686243, KJ686186, KJ686154, HQ129932, KJ686257, KJ686255, KJ686228, KR045297, DQ341356, JN864022, KJ686169, MG756718, HQ647167, KY014078, KJ686288, KJ686271, KJ686227, KJ686207, MF662683, KJ686275, MG756717, MG756715, KJ686291, KJ686242, JF894382, KJ686282, KJ686281, KJ686276, KJ686235, KJ686215, KJ686205, KJ686150, KJ686145, KP308431, MG756716, KT428650, KJ686164, KJ686253, KJ686146, HQ647174, KJ686240, KJ686233, KJ686183, KJ686139, HQ647176, KJ686285, MF662699, KJ686173, KJ686136, MF662700, MG976582, KJ686209, HQ647168, MG756691, KJ686167, KJ686138, KU936131, KJ686304, KJ686190, MG756692, LC321992, KP308414, KJ686301, KP308458, KJ686247, KR045304, KJ686294, KJ686232, KR045293, KP308425, GU198371, KP308429, KJ686260, GU198370, KJ686278, KJ686212, KX197455, KP308418, KJ686293, KR045292, KR045291, KP289426, KP289420, KP308445, KX197461, KP308424, KM211579, KU936129, KP308421, KT428644, KR045294, KX197456, KR045295, MF405075, KT428648, MG756719, JF820315, KP308446, HQ647169, KP308444, JF820312, HQ424434, HQ424435, KC296445, KP274876, JF820316, JF820313, MG773124, MG207963, FJ172159, HM752814, JF820314, AB575941, KC296443, HQ423143, HQ424433, KX430820, KP308406, KX430823, KP274874, KP274875, KP308417, KR045303, KX430819, KM593929, KJ686187, KJ686144, KJ686191, KJ686179, KJ686230, AF119795, EU527983, KP308456, KP308411, KJ632499, KP308416, KP308448, KX430818, LC321993, MH718269, KP308427, KF543271, KX197465, KP308404, KP308447, KP308420, KF306100, HM156065, KU159440, KX588236, KU159436, KP308435, KJ186974, KF306101, HQ850973, JN230523, KU159435, KU159441, MH511208, MH511205, MH511194, MH511182, MH511207, MH511202, MH511198, MH511197, MH511196, MH511195, MH511191, KU159442, MH511204, MH511192, MH511187, MH511185, MH511184, MH511181, KU159439, MH511206, MH511203, MH511201, MH511200, MH511199, MH511190, MH511188, MH511186, GU350629, MH511193, JQ950555, KU159437, KU159438, KJ686134, KU159434, KX430817, AB747374, MG013988, MG672480, MK800119, KF974786, AB575916, LC321991, FJ357379, FJ357383, MH716378, KX372315, KF974782, MG367596, KT354873, FJ357381, AB550336, KY888026, KF974788, KF514879, MG756695, MH716263, FJ357378, MG756707, JN964686, MH716392, DQ341363, MH716262, KJ686296, KX372329, MH716382, MH716307, MG756754, MH716359, MH716363, MH716306, KP266579, MG756694, MH716312, AB550332, KP691644, MH716365, KP691649, KJ746493, MH716289, MH716301, FJ607334, MH716371, JX678874, DQ341366, KX372325, KU647000, KX372312, DQ452074, LC375766, JN052925, KC436267, FJ357377, MH716343, KP691661, KC954662, MH716372, AF304457, MH716368, MH716345, DQ341361, EF373575, EU753397, GQ231933, FJ360544, HQ456309, HM053670, MH716320, MH716357, MH716322, AB550340, KX372321, GQ231938, AB575915, MH716295, FJ357373, MF662682, GQ994990, MH716367, KF142412, EU753384, MH716375, MH716326, JN864020, JX244182, KF312457, MH716338, JQ319054, GU366191, DQ133459, JN256062, KP691646, FJ461781, MH716330, MH373639, MG208882, JN992283, HM002484, MH716264, JN001860, KC436268, KP266572, JQ086365, KJ004558, KF444809, KX197464, DQ133458, HQ456312, AB575912, FJ606449, JN256061, MH716339, JQ806378, KP289421, HQ456308, MH716249, MF662688, KC436272, JQ514785, MG756701, MG756703, MK344771, KJ686254, KX372319, HQ998852, DQ341354, KP308443, JX111889, HM002488, GQ892830, HQ456307, MF662679, MG756724, JX244187, JN864019, KJ004557, KJ686292, HQ825317, HQ456313, KC414134, HQ400942, KJ686244, JQ804832, JX986737, HQ407557, JF738000, KM402021, KJ686265, KM591215, HQ325852, HQ188292, KJ004554, MF662690, KP198623, KJ686272, KP198624, KJ784496, KJ004555, JQ074190, EF373576, MG212485, JQ736684, JQ086366, KJ686218, MF662693, KT428645, HQ423142, MG773123, KX372309, MG756720, KJ004552, HQ828086, KP289428, HM245927, MF362981, KU936124, KP289417, MF662677, KJ004553, MF662697, MF662689, HQ712020, KX197460, KJ686220, KC570452, HQ891929, HQ426649, MF662691, KJ686177, KJ686219, MF662696, KP289423, KJ686148, KP308405, KP861243, JN864023, MF662694, HQ647175, HQ891926, MK307505, KU936123, KJ686199, MF662692, KJ686203, KJ686147, KJ508182, HM245928, KJ686133, AB575937, KU254595, KC570453, HQ647170, FJ828519, KJ686290, KP289429, MG756693, KJ686274, KJ686273, KT345960, KP289430, KF668443, KU936126, MG875331, KP289427, KJ686180, KY582572, KJ686250, KM211580, MG520666, KU936127, JN835312, MF431793, KU936122, KR045296, KJ686135, MG581490, HQ424436, JQ639383, KF142411, EU812515, MG773122, KP289418, MF662698, AB575942, AB575948, HQ424437, AB575939, MG756705, KX893543, GU196833, KY888025, KJ632497, KJ686178, KJ632498, MH511183, MH511189

**Porcine KoV	GenBank	All**

MF506730, AB624493, LC210613, LT898428, GQ249161, EU787450, LC210612, LC210606, LC210600, LC210601, LC210603, LC210602, LC210610, LC210604, LC210620, LC210599, JQ692069, MG800803, MG800804, MG800807, MG800806, LC210611, LC210622, MF062451, MF062445, MF062449, LC210618, KC424640, KC424638, KC424639, KY234500, GU292559, LC210617, LC210616, MK962328, MK962332, MK962335, MK962336, MK962321, MK962330, MK962320, MK962333, MK962334, MK962323, MK962327, MK962329, MK962331, MK962324, JX177612, MK962322, MK962326, MK962325, LC210608, LC210609, LC210605, LC210607, LC210619, LC210621, KM977675, KP144318, KP260507, KT892971, LC210614, LC210615, JN630514, JX827598, MF062440, KJ452348, MF062448, MF062438, MF062437, KC204684, MG800805, JX401523, KM051987, MF062436, KC414936, KF539763, KF695124, MF062447, MF062443, MF062444, MF062450, MF062452, MF062441, MF062442, MF062446, MF062439, KY234499, MF062435

**DENV1	GenBank	All**

AB608788, AB608786, AB608787, LC016760, JX669468, KF971870, KF971871, GU131922, EU848545, KC172829, AF298808, FJ196847, KF955441, KF921948, AF514889, EU482567, AF180817, JN903580, JQ922544, GU131887, AY277666, AY722802, AY708047, JQ922545, denv, LC011948, AB189121, AB195673, AB204803, KF921911, KC762631, KC762652, AB074760, JX669463, JX669465, HM469967, JN903579, FJ850070, GU131956, KF955442, GU131966, KJ189349, KJ189350, FJ687426, FJ205875, FJ410188, FJ850099, DQ285560, EU081257, KF955406, AY732483, KP723476, AF350498, KF672763, FJ196846, AB178040, KJ468234, HE795086, LC011945, LC011949, HM469966, JX669462, JX669464, JX669466, JX669467, JX669469, JX669470, JX669471, JX669472, JX669473, JX669474, JX669475, KF971869, JN903578, JN903581, KJ649286, KF887994, FJ850073, FJ850075, FJ850077, FJ850081, FJ850084, FJ850087, FJ850090, FJ850093, GU131863, GQ868559, GQ868560, GQ868561, GU131948, GU131949, GU131888, GU131889, GU131890, GQ868630, GU131891, GU131892, GU131893, GU131894, GU131895, GQ868632, GQ868633, GU131919, GU131920, GU131921, GQ868635, GQ868636, GU131923, GQ868637, GQ868639, GU131925, GU131926, FJ639669, FJ639670, FJ639671, JN819423, FJ639672, FJ639673, FJ639674, FJ639675, FJ639676, FJ639677, GQ868618, GQ868619, FJ639678, FJ639679, FJ850069, FJ639680, FJ639682, JQ287664, FJ639683, FJ639684, FJ639685, FJ639686, FJ744702, FJ639688, FJ639689, FJ639690, FJ639691, FJ639692, FJ639693, FJ639694, FJ639695, FJ639696, HM631852, KF955444, HM181936, HM181937, HM181938, HM181939, HM181940, HM181941, HM181942, HM181943, HM488255, HM181944, HM181946, JQ287665, HM181948, HM181949, HM181950, HM181951, HM181952, HM181953, KF921933, HM181954, HM181955, HM181958, HM181959, JF937651, KF955415, FJ687428, FJ687429, FJ850068, FJ687430, FJ687432, FJ687433, JQ922547, EU660390, EU660391, EU660392, EU660393, EU660394, EU660401, FJ373305, EU660402, EU660403, EU660397, EU687247, EU660395, EU660412, EU660418, EU677150, EU677151, EU677152, EU677153, EU677154, EU677155, EU677156, EU677157, EU726777, EU677158, EU677159, EU677160, EU677161, EU677162, EU677163, EU677164, EU677165, EU677139, EU677140, EU677166, EU677167, EU677168, EU677170, EU726778, EU677171, EU677172, EU677173, EU677174, EU726779, EU660419, EU677175, EU677176, EU726780, EU726781, EU677177, EU726782, EU677178, EU687251, FJ024426, FJ024427, FJ024428, FJ024429, FJ024430, FJ024431, FJ024432, FJ024433, FJ024434, FJ024435, FJ024436, FJ024437, FJ024472, FJ024439, FJ024441, FJ024442, FJ024443, FJ024444, FJ373296, FJ182018, FJ410287, FJ182019, FJ432719, FJ182020, FJ024445, FJ182021, FJ182022, FJ024446, FJ182023, FJ024447, FJ182025, FJ373297, FJ024448, FJ182026, FJ373298, FJ182027, FJ390381, FJ182028, FJ024449, FJ390382, FJ182003, FJ182029, FJ390383, FJ024450, FJ182030, FJ182031, FJ182033, FJ182034, FJ182036, FJ410289, FJ205876, FJ024455, FJ024456, FJ024457, FJ390386, FJ024459, FJ205881, FJ024460, FJ205882, FJ205883, FJ024462, FJ024463, FJ390388, FJ024464, FJ410191, FJ432723, FJ410192, FJ410194, FJ432725, FJ432727, FJ432729, FJ432730, FJ432732, FJ432733, FJ410196, FJ410197, FJ432734, FJ432735, FJ432736, FJ410198, FJ432737, FJ410199, FJ432738, FJ432739, FJ432740, FJ410201, FJ410203, FJ410204, FJ432742, FJ410205, FJ410206, FJ410207, FJ432744, FJ547060, FJ432745, FJ410209, FJ410210, FJ432746, FJ410211, FJ432747, FJ432748, FJ432749, FJ461306, FJ410212, FJ461307, FJ410213, FJ461308, FJ461310, FJ410216, FJ461312, FJ906728, FJ461313, FJ547063, FJ410218, FJ461315, FJ461316, FJ461317, FJ410220, JF937649, FJ410222, FJ461318, FJ461319, FJ461320, FJ410225, FJ410226, FJ410227, FJ859029, FJ547065, FJ410230, FJ410231, FJ461323, FJ461324, FJ461325, FJ461327, FJ461328, FJ410234, FJ410235, FJ410236, FJ410238, FJ410239, FJ410240, FJ410242, FJ410243, FJ410244, FJ410245, FJ461330, FJ410246, FJ410247, FJ410248, FJ410249, FJ410250, FJ410251, FJ410252, FJ410254, FJ461331, FJ410255, FJ410256, FJ410257, FJ461332, FJ410258, FJ461333, FJ410260, JF937650, FJ410261, FJ461335, FJ410262, FJ410263, FJ410264, FJ410265, FJ461336, FJ410266, FJ410267, FJ410268, FJ410269, FJ410270, FJ410272, FJ410273, FJ410274, FJ410275, FJ410276, FJ410277, FJ410278, JQ287667, FJ410279, FJ410280, FJ410281, FJ410282, FJ410283, FJ410284, FJ410285, FJ410286, FJ461339, FJ461340, FJ461341, FJ882515, FJ882516, FJ882517, GQ199771, KF921934, JQ287660, JQ287661, FJ882518, FJ882519, FJ882520, FJ882521, FJ882522, FJ882523, FJ882524, FJ882525, FJ882526, FJ882527, FJ882528, FJ882529, FJ882530, FJ882531, FJ882532, GQ868605, FJ882533, FJ882534, FJ882535, FJ882536, FJ882537, GQ868606, FJ882538, FJ882539, FJ882541, GQ199772, FJ882542, FJ882543, FJ882544, FJ898371, FJ882545, FJ898372, FJ898373, FJ898374, FJ898375, FJ898376, FJ898377, FJ898378, FJ898379, GQ199773, GQ868607, JF937596, FJ906963, FJ906964, FJ906965, FJ898380, FJ898381, FJ898382, FJ898383, FJ898384, FJ898385, GQ868608, GQ199774, GQ199775, JQ287662, FJ882546, KF921935, GQ199776, GQ199777, JQ287663, GQ199778, GQ199779, GQ199780, GQ199781, GQ199782, GQ199783, GQ199784, GQ199785, GQ199786, GQ199787, GQ199788, GQ199789, GQ199790, GQ199791, GQ199792, GQ199793, GQ199794, GQ199795, FJ882547, GQ199796, GQ199797, GQ199798, GQ199799, GQ199800, FJ882548, GQ199801, GQ199802, JF937597, GQ868609, GQ199803, GQ199804, GQ199805, GQ199806, GQ199807, GQ199808, GQ199809, GQ199810, GQ199811, GQ199812, GQ199813, FJ882549, GQ199814, GQ199815, GQ199816, GQ199817, GQ199818, FJ882550, FJ882551, GQ199819, FJ882552, FJ882553, FJ882554, GQ868610, GQ868611, GQ199820, FJ882555, GQ199821, FJ882556, GQ199822, FJ882557, FJ882558, GQ199823, GQ199824, FJ882559, FJ882560, GQ199826, GQ199827, GQ199828, FJ882561, GQ199829, FJ882562, FJ882563, FJ882564, FJ882565, GQ199830, FJ882566, FJ882567, GQ199831, GQ199832, FJ882568, FJ882569, GQ199833, FJ882570, GQ199834, GQ199835, FJ898386, GQ199836, GQ199837, FJ898387, FJ898388, FJ898389, FJ898390, FJ898391, FJ898392, FJ898393, FJ898394, FJ898395, GQ199838, FJ898396, GQ199839, FJ898397, FJ898398, FJ898399, FJ898400, FJ898401, FJ898402, FJ898403, FJ898404, GQ199840, FJ898405, FJ898406, GQ199841, GQ199842, FJ898407, FJ898408, GQ199843, FJ898409, GQ199844, GQ199845, JF937598, GQ199846, GQ199847, GQ199849, GQ199850, FJ898410, GQ199851, FJ898412, GQ199852, FJ898413, FJ898414, FJ898415, GQ199853, GQ199854, FJ898416, FJ898417, GQ868612, GQ199855, FJ898418, FJ898419, FJ898420, FJ898421, FJ898422, FJ898423, GQ868613, FJ898425, FJ898426, GQ199856, FJ898427, FJ898428, FJ898429, JF937599, FJ898431, GU131678, GU131679, GU131680, GU131681, GU131682, GU131683, GU131684, GU131685, GQ868614, GU131686, GU131687, GU131688, GU131689, GU131690, GU131691, GU131692, GU131693, GU131694, GU131695, GU131696, GU131697, GU131698, GU131700, GU131701, GU131702, GU131703, GU131704, GU131705, GU131706, GU131708, GU131709, GU131710, GU131711, GU131712, GU131713, GU131714, GU131715, GU131716, GU131717, GU131718, GU131719, GU131720, HM181960, HM181961, HM181963, HM181964, HM181965, HM181966, GU131721, GU131722, GU131723, GU131724, GU131725, GU131726, GU131727, HM631850, GU131728, KF955446, KF921942, JF937602, GU131729, GU131730, GU131731, GU131732, GU131733, GU131734, GU131735, GU131736, GU131737, JF937603, GU131738, JN000935, JF937604, JF937605, JF937606, JF937607, JF937608, GU131739, GU131740, GU131741, GU131742, GU131743, GU131744, GU131745, JN093516, GU131746, JF937609, KF921949, GU131747, GU131748, GQ868615, GU131749, GU131750, GU131751, GU131752, GU131753, GU131754, GU131755, GU131756, GU131757, GU131758, GU131759, GU131760, GU131761, GU131762, GU131763, GU131764, GU131765, GU131766, JF937610, GU131767, HM181967, JF937611, GU131768, GU131769, GU131770, GU131771, GU131772, GU131773, GU131774, GU131775, GU131776, GU131777, GU131778, GU131779, GU131780, GU131781, GU131782, GU131783, GU131784, GU131785, JF937612, GU131786, GU131787, GU131788, HM181968, GU131789, GU131790, HM181969, GU131791, GU131792, HM488256, GU131793, GU131794, GU131795, GU131796, GU131797, GU131798, GU131799, GU131800, GU131801, JF937613, JF937614, GU131802, GU131803, GU131804, GU131805, HM631851, GU131806, GU131807, GU131808, GU131809, GU131810, GU131811, JF937615, GU131812, GU131813, GU131814, GU131815, GU131816, GU131817, GU131818, GU131819, GU131820, HM181970, GU131821, GU131822, GU131823, GU131824, GU131825, GU131826, JF937616, JF937617, JF937618, JF937619, GU131827, GU131828, GU131829, GU131830, GU131831, EU482789, EU482790, EU482791, EU482792, EU482706, EU482707, EU482708, EU482709, EU482710, EU482711, EU482712, EU482713, EU482714, EU482715, EU482717, EU482718, EU249490, EU249491, EU249492, EU249493, EU249494, EU249495, EU482793, EU482795, EU482796, EU482797, EU482798, EU482799, EU482800, EU482802, EU482803, EU482804, EU482805, EU482806, EU482807, EU482808, EU482809, EU482810, EU482811, EU482812, EU482813, EU482814, EU482815, EU482816, EU482817, EU482818, EU482819, EU482820, EU482821, EU482822, EU482823, EU482824, EU482825, EU482826, EU482827, EU482828, EU482476, EU482477, EU482478, EU482479, EU482481, EU482482, EU482483, EU482484, EU482485, EU482486, EU482487, EU482488, EU482489, EU482490, EU482491, EU482492, EU482493, EU482494, EU482495, EU482496, EU482497, EU482498, EU482499, EU482500, EU482501, EU482502, EU482504, EU482505, EU482506, EU482508, EU482509, EU482510, EU482511, EU482512, EU482513, EU482514, EU482515, EU482516, EU482517, EU482519, EU482520, EU482521, EU482522, EU482523, EU482524, EU482525, EU482526, EU482527, EU482528, EU482532, EU482533, EU482539, EU482540, JQ045626, JQ045628, JQ045629, JQ045630, JQ045631, JQ045632, JQ045633, JQ045634, JQ045636, JQ045635, JQ045637, JQ045638, JQ045639, JQ045640, JQ045641, JQ045645, JQ045642, JQ045644, JQ045646, JQ045647, JQ045648, JQ045649, JQ045650, JQ045651, JQ045652, JQ045653, JQ045654, JQ045656, JQ045657, JQ045658, JQ045659, JQ045660, JQ045661, JQ045662, JQ045663, JQ045664, JQ045665, JQ045666, JQ045667, HQ624983, FJ384655, JQ048541, JN697056, JN697057, HQ624984, KP398852, HQ891313, HQ891314, HQ891315, HQ891316, JN054256, JN054255, KJ438293, KJ438296, KC692495, KC762625, KC762654, KC762636, KC762635, KC762650, KC762623, KC762632, KC762646, KC762637, KC762620, KC762621, KC762643, KC762651, KC762638, KC762640, KC762628, KC762644, KC762630, KC762641, KC762648, KC762622, KC762634, KC762633, KC762629, KC762627, KC762647, KC762639, KC762642, JQ915077, GU370049, HM469968, KC172834, KJ933413, EU359008, AY835999, KF672760, FJ176780, GQ398255, JN205310, HQ332182, KP406803, KP406801, KC131140, KT279761, KP723473, KJ726663, KJ726662, KJ189316, HM631855, JN638344, JN638343, JN638342, JN638341, JN638340, JN638339, JN638338, JN638337, JN638336, FJ469909, FJ469908, FJ469907, AB519681, EF025110, AF226685, AY145123, EU081254, EU081281, EU081280, EU081279, EU081278, EU081277, EU081276, EU081275, EU081274, EU081273, EU081272, EU081271, EU081270, EU081268, EU081267, EU081266, EU081265, EU081264, EU081263, EU081260, EU081259, EU081258, EU081256, EU081255, EU081253, EU081252, EU081251, EU081249, EU081248, EU081247, EU081246, EU081245, EU081244, EU081243, EU081241, EU081239, EU081238, EU081237, EU081236, EU081235, EU081234, EU081233, EU081232, EU081231, EU081230, EU081229, EU081228, EU081227, EU081226, EF457905, AY732482, AY732481, AY732480, AY732479, AY732478, AY732477, AY732476, AY732474, HG316481, U88537, AB074761, AF298807, AY726555, AY726554, AY726553, AY726552, AY726551, AY726550, AY726549, AY722803, AY713475, AY713474, AY713473, KF864667, JQ915078, FJ196843, FJ196842, KJ726665, KF921951, KF921943, KF921940, KF921939, KF921936, KF921932, JQ692085, JQ917404, GU370048, DQ285558, KP686070, KJ726664, AY732475, AY762084, EU179861, AY722801, AY713476, AB189120, GQ868602, JN697058, KC762653, KC762645, KC762624, KC762626, JQ915079, JQ915080, JQ915072, JQ915071, JQ915073, JQ915074, JQ915075, JQ915076, KF289072, KC172835, FJ176779, KM204119, DQ285562, DQ285559, AF311956, KP406802, DQ672563, DQ672562, DQ672560, DQ672559, DQ672558, DQ672557, DQ672556, AF309641, JQ045627, AY145122, AY145121, AF226687, EF032590, HG316482, U88535, AF226686, FJ196848, FJ196845, FJ196844, FJ196841, KR028435, JX669461, KF184975, JQ675358, FJ850071, GQ868562, GQ868563, GQ868564, GQ868565, GQ868566, GQ868567, GQ868568, GQ868569, GQ868570, KJ189302, KJ189303, KJ189304, JQ922548, JQ922546, GU131957, GU131958, GQ868498, KF955416, KF955417, GQ868499, GQ868500, GU131960, GU131961, GU131962, GU131963, GU131964, KF955419, GU131965, KF955421, KF955422, GQ868501, HQ166035, GQ868502, GQ868503, GQ868504, GU131967, GQ868505, GU131968, GU131969, GQ868506, GQ868507, GQ868508, GQ868509, GQ868510, GQ868511, GQ868512, GU131970, KF955427, GU131971, GQ868513, GQ868514, GU131972, GU131973, GQ868517, GU131976, GQ868518, GQ868519, KF955443, GQ868520, GQ868521, GQ868522, GU131977, GQ868523, GQ868524, GU131978, HQ166036, GQ868525, GQ868526, GU131979, GQ868527, GU131980, GU131981, GQ868528, GQ868529, GU131982, GQ868530, GQ868531, GU131983, GQ868532, KF955433, GQ868533, GQ868534, GQ868535, KJ189306, KJ189307, KJ189318, KJ189319, KJ189320, KJ189321, KJ189322, KJ189324, KJ189325, KJ189326, KJ189327, KJ189328, KJ189329, KJ189330, KJ189331, KJ189332, KJ189333, KJ189334, KJ189335, KJ189336, KJ189337, KJ189338, KJ189339, KJ189340, KJ189341, KJ189342, KJ189343, KJ189344, KJ189345, KJ189346, KJ189347, KJ189348, KJ189368, KJ189369, EU482615, EU482616, EU482617, EU482618, EU482619, FJ898433, FJ547088, FJ810419, FJ547068, FJ562104, GQ199867, GQ199857, GQ199858, GQ199859, JF937644, JQ287666, JF937645, FJ850114, JN819402, FJ850113, JF937635, FJ024483, EU596504, FJ024481, GQ199872, FJ410290, FJ024423, FJ024485, FJ547089, FJ024484, EU596503, FJ024479, FJ182002, FJ024482, EU596501, FJ024480, GQ199873, FJ873814, GQ199875, JN819403, KF973453, KF973454, KF973455, KF973456, KF973457, KF973458, KF973459, KF973460, KF973461, KF973462, KF973463, KF973464, KF973465, KF973466, KF973467, KF973468, KF973469, KF973471, KF973472, KF973473, KF973474, KF973475, FJ898448, KJ189351, KJ189352, KJ189353, KJ189354, KJ189355, KJ189356, KJ189357, KJ189358, KJ189359, KJ189360, KJ189361, KJ189362, KJ189363, KJ189364, KJ189365, KJ189366, KJ189367, JN819417, FJ390374, FJ390378, FJ205872, FJ205873, FJ390379, FJ390380, FJ205874, FJ410173, FJ410174, FJ410175, FJ562105, FJ562106, FJ410179, FJ410181, FJ547086, FJ410183, FJ410184, FJ410185, FJ547087, FJ410186, FJ478457, FJ410189, FJ478458, FJ410190, EU482591, EU482592, EU482609, EU482610, EU482611, FJ639735, FJ639740, FJ639741, FJ639743, FJ639794, FJ639796, FJ639797, FJ639802, FJ744701, JN819410, FJ639806, FJ639808, FJ639811, JN819411, FJ639812, FJ639813, JN819412, FJ810415, FJ639814, FJ639815, FJ639818, FJ639819, FJ639820, FJ639821, FJ639823, FJ639824, JN819413, JN819425, KF955407, FJ850100, FJ882579, JN819414, FJ873810, FJ850103, FJ850104, JN819405, GQ199877, GU056029, GU056030, GU056031, GU056032, GU056033, GU131833, GU131834, GU131835, GU131836, GU131837, JN819415, GU131838, GU131839, GU131840, GU131841, GQ868601, EU660396, KC759167, KC692496, KC692497, KC692498, KC692499, KC692500, KC692501, KC692502, KC692503, KC692504, KC692505, KC692506, KC692507, KC692508, KC692509, KC692510, KC692511, KC692512, KC692513, KC692514, KC692515, KC692516, KC692517, KC762649, KF672759, KF672761, KF672762, JF459993, KF672764, EU863650, HQ332177, HQ332181, HQ332180, HQ332183, HQ332179, DQ285561, AF311958, AF311957, EU280167, AY277665, AY277664, AY277659, KC131141, KF289073, KJ189315, KJ189314, KJ189313, KJ189312, HQ166037, HM631853, GU131984, GQ868539, GQ868537, GQ868536, FJ461303, EF122232, EF122231, DQ672564, AF513110, DQ193572, EU081262, M87512, KF955428, KF955420, AY206457, AF514883, AF514885, AF514876, AF514878, AF180818

**HEV	VIPR database	Selection**

MH992007, MF444058, MF444057, MH377722, MH504140, MH809516, MH504142, AB193178, MF444040, MF444073, MH992008, AB220971, MH992001, KC492825, LC314156, MH504151, JF443726, MH504148, MN646691, MF346772, MH504157, MF444138, MF444030, MH504150, MF444047, MF444117, AB437316, MF444048, MF444077, MH992010, AB080575, HQ389543, KC618403, LC037955, MF444104, MF444064, MF444097, LC022745, JQ679013, MF444096, MH377724, MN401237, AB521805, KU176131, MH504163, MH504132, MF444130, MH377725, AB437317, KY436507, FJ653660, JQ013795, AB193177, MF444035, MF444087, AB074920, MF444109, MF444084, MF444050, MF444108, MF444106, MH504133, LC387631, AP003430, MH504149, MF444088, MF444105, MK089849, AB097812, EU495148, JQ679014, KR872415, MF444072, JQ655734, MF444041, MF444056, MF444082, HM439284, MF444051, MF444054, MF444029, MF444093, MF444055, MF444061, MF444079, MF444037, MF444123, MH504143, MH991996, MF444033, MF444125, MF444067, MF444042, MF444101, MH504127, MH377721, X98292, HQ634346, MF444043, MF444134

**HNoV, genogroup II	VIPR database	Selection**

KJ196282, KP784694, KJ196287, AB933700, AY502020, AB933677, JN400613, MG049692, KJ196291, AB933676, GQ845024, MF405169, KU561248, KP784691, KU561252, NC_044045, KP784693, KJ196294, KT970369, LN854572, KJ196278, KJ196296, KJ196288, KJ196281, KJ196283, KJ196299, KT970375, KP784692, KT970373, KT380915, KT970372, LN854565, KP784695, KU561251, KJ196295, KJ196293, KT202797, KJ196280, KT202796, KJ649705, KJ196276, LN854570, NC_039475, KP784696, KU561254, KX079488, KP998539, LN854566, KU561249, KT202795, KU870455, KT970377, KU561250, KJ196277, KT202794, LN854569, KJ196285, LN854568, KU561253, NC_044046, KJ196290, LN854567, KU561255, KT970374, AY502023, KU561256, KJ196297, KT202793, KJ196289, KT202798, KT970371, KP784697, KT589391, KJ196284, KP784698, LN854571, KT970376, KT970370, KJ196279, KJ196286, KF429790, AB933648, KY865306, LC145787, MK762560, MG746035, MK775032, KM198573, MK282256, KF429785, MK762633, KF712499, KY947549, GU980585, MH218597, MK754442, KC409272, KC409249, KJ685413, MK775029

**JEV	GenBank	All**

JN381838, KT957421, JN381872, JN381830, JQ086763, JN381834, AF075723, HQ652538, AB241119, JN381867, AY184212, AB551990, AB569988, AF217620, KM658163, AB471669, AY303796, AB051292, KY927815, KF297916, HQ223286, M18370, JF706283, JN381853, AF080251, KY927818, AY316157, GQ902063, JF915894, KY078829, JN381863, AB551991, AF045551, KT229572, L48961, HQ223287, JN381843, JF706270, HQ223285, GQ902062, GQ902061, KM677246, KT957420, AY303794, KT229575, JF706279, HE861351, JN381868, JN381870, KY927816, U47032, AF098736, EF623987, JN381873, AB241118, JN381848, JN381866, AF221500, AY303792, GU205163, KF667311, GQ902060

**HPgV-1	GenBank	All**

AF121950, D87255, AF081782, D90600, LT009483, LT009487, LT009485, LT009494, AY196904, MH053119, LT009481, LT009478, HGU44402, LT009479, AF309966, MK291245, MK291244, AF031827, MH053115, KP259281, MH053120, HGU45966, MH053118, MH053121, HGU63715, LT009486, AB003289, LT009489, AF104403, JN127373, D90601, AB013501, MH053116, D87711, HGU36380, AB008335, D87263, D87712, D87713, KM670109, D87262, D87709, KM670096, LT009490, AB008342, KM670099, D87714, LT009480, KP710601, KM670110, LT009482, D87710, AB021287, KM670097, LT009484, KP710599, HQ331235, KM670100, HQ331234, LT009488, KM670107, HQ331233, KM670108, KP710604, KM670102, KM670106, KC618400, D87708, AB008336, KM670098, HGU94695, KP710605, KP710600, KC618401, MH746815, KC618399, AB013500, KP710598, KC618398, KM670101, MH053117, KP710602, KP710603, AB003288, KP710606, AB018667, HGU75356, AY949771, D87715, AB003290, AB003293, AF006500, AB003291, AB003292, MH179063, LT009493, MK291243, LT009492, LT009491, AF121950

**MNV	GenBank	All**

FJ446719, JN975495, EU004682, DQ223042, EU004668, AB435515, EU004680, JF320653, JN975498, EU004675, JF320647, DQ223043, EU004671, FJ446720, EU004674, EU004670, EU004681, EU004658, EU004677, HQ317203, EU004667, JN975493, JF320652, EU854589, JN975496, EF531290, AB601769, JN975494, JQ237823, EU004678, EU004663, DQ911368, EF531291, EU004683, EU004679, EU004664, DQ223041, EU004673, DQ285629, EU004656, EU004659, EU004661, EU004662, AY228235, EU004655, EF014462, EU004654, EU004657, JF320648, JF320649, JF320650, JF320645, JF320651, JF320646, EU004669, EU004665, EU004672, EU004660, AB435514, JF320644, EU004676, JQ658375, EU004666

**Canine KoV	GenBank	All**

KF924623, MN449341, KC161964, MH052678, JN387133, JN088541, MF062158, JQ911763, MK201776, MH747478, MK201779, MK201777, MK201778, KM068051, KM068048, KM068050, KM068049, KF831027, MF598159, KM091960, MK671314, MK671315, MH159813, MH159814, KJ958930

**FMDV-A	GenBank	All**

JF749843, AY593766, AY593770, AY593756, AY593793, AY593788, AY593782, AY593758, AY593760, AY593771, AY593752, AY593759, GQ406251, HQ832577, HM854024, HQ832576, HQ832585, KJ754939, HQ832591, HQ832584, HQ832589, AY593761, AY593794, AY593768, AY593775, AY593803, AY593790, AY593789, AY593792, AY593767, AY593776, AY593779, AY593755, JN099699, EF494487, JN006722, KM268896, KJ933864, AY593765, HM854022, JF749848, HQ832586, HQ832592, HQ832580, HQ832587, AY593773, AY593787, AY593786, AY593802, AY593801, AY593785, AY593784, AY593783, AY593769, AY593753, AY593757, AY593781, AY593780, AY593778, AY593754, APHA12CDR, AY593777, AY593774, AY593751, JN099698, JN099697, JN099688, JN099695, JN099694, EF494488, EF117837, JF749841, EF494486, KC588943, GQ406250, GQ406249, GQ406252, GQ406248, GQ406247, HQ268509, HQ632773, HM854023, AY593764, AY593763, AY593762, X74812, AY593772, AY593791, HQ832590, HQ832583, HQ832579, HQ832578, HQ832581, HQ832582, HM854025, HM854021, HQ832588,

**FMDV-O	GenBank	All**

AY593824, JN998086, FJ461345, DQ248888, AY686687, EF552691, JX066664, JX040492, KF694737, AY593813, AY593822, KJ206910, KJ825804, AY593826, KF112882, HM191257, GU384683, HM008917, EU448369, AY593817, GU125647, AY593833, KF112880, KF112885, JX869177, HQ632769, AY593827, AF511039, AY333431, DQ478937, KF112888, FJ461344, AY593812, HQ412603, HQ113232, KJ206908, HQ632768, AY359854, FJ175664, KF112879, JQ900581, AY593830, AY317098, AY312588S2, GU125650, KF501486, HQ009509, AY593834, AF377945, DQ119643, KF112886, EU140964, HQ632770, KF501488, KM268895, HQ632771, JX869178, AY593825, JF749851, HQ632772, HQ268524, AY593823, AY593821, KF985189, KJ206909, JX040501, AY593828, AY593829, KJ825801, AB079061, AY593814, JX040488, AY312586S2, DQ404179, JX040497, EF552696, JX040493, JN998085, KR265072, JX040490, FJ542366, DQ404169, DQ404170, AJ320488, FJ542370, EU214601, JX040491, DQ404164, HM055510, KR265074, JX570643, EF614457, DQ404166, EF552695, DQ404162, JX869186, JX570653, JX570652, DQ404168, AY593820, KF694740, DQ404165, EF552690, EF552694, GU125648, KJ825808, AY593815, JX869179, KF694731, JX869182, AY593818, JX570644, JX869181, EU448381, JX570654, EU448379, JX869185, DQ404175, DQ404167, AY593811, JX869180, JX570655, AF154271, DQ404159, EU448376, FJ175662, JX040495, EF552692, AJ633821, EU448377, AJ539140, JX869184, EF552697, JX570647, KR265075, KM257062, GU384682, FJ542369, KM257063, JX570641, KF694733, FJ175665, KJ825806, AJ539137, KJ825807, JX040487, AJ539139, EF552689, KM257065, KF112887, JX947858, DQ478936, EU448373, KF112881, EU448372, DQ404174, DQ404160, JX570645, DQ404176, JX570640, JX869187, EU448368, KC503937, AY593816, KJ825803, KF694743, AY593836, EU448378, JX570638, EU448375, JX040499, JX869188, KJ825805, KF112883, FJ542365, KF694745, KM257064, JX570648, JX570650, EU448371, AF506822, FJ542372, KF694739, KF112884, FJ175661, EF175732, EU448380, JX040485, DQ404173, FJ175663, JX040489, X00871, DQ404161, EU448370, FJ542371, DQ404172, KF694735, JX040498, EF552688, JX040494, EF552693, FJ542368, JX570651, DQ404158, DQ404177, AY593835, JX040486, KF694741, KJ825809, KR401161, AF308157, JX570649, FJ175666, JQ973889, DQ404180, JX869183, JX040496, KF112889, KF694744, KF694736, EU448374, KJ825802, JX570639, KR265073, AY593831, AY593837, DQ404163, AJ539136, GU125649, JX066665, AY593819, AY593832, KF694742, KM257061, KF694732, KF694738, KF501487, DQ404178, DQ404171, AJ539138, AJ539141, JX570642, JX040500, JX570646, FJ542367, HM229661

**RUBV	VIPR database	All**

AY258323, AY258322, AB928203, AB928205, AB928204, KX291007, FJ211588, FJ211587, CS406445, AB588192, AB047330, AB047329, AB222608, KT962867, KT962871, JN635284, JN635287, KT962864, KT962869, KT962870, JN635288, JN635281, AB222609, JN635283, AB860305, KT962866, KF201674, JN635291, KT962863, JN635285, JN635282, JN635286, L78917, JF727654, JF727653, AB588190, AB588191, AB588189, DQ388281, NC_001545, M15240, DQ085339, AX009468, AF188704, DQ085340, DQ388279, DQ085341, DQ085343, MF496142, AB588188, KT962868, JQ624625, JN635293, JN635294, JN635292, KT962865, JN635295, JQ624624, JN635296, KT962862, AB588193, DQ085338, DQ085342, JN635290, JN635289, DQ388280, X72393, AF435865, AF435866, KU958641, KT000088, KT000089, AF533117

TABLE S2

COMPOSITIONAL FEATURES OF RNA VIRUS SEQUENCE DATASETS USED IN THE STUDY

|  |  |  | |  |  |  | **Base imbalance^3^** | | **Dinucleotide Rep.^4^** | |  |  | **Transition Asymm^5^** | | **Normalised Asymm^6^** | |
| --- | --- | --- | --- | --- | --- | --- | --- | --- | --- | --- | --- | --- | --- | --- | --- | --- |
| **Virus^1^** | **Family** | | **Polarity** | **n** | **MPD^2^** | **G+C** | **C/G Asymm** | **U/A** | **CpG** | **UpA** | **MFE** | **MFED** | **rG->A** | **rC->U** | **nG->A** | **nG->U** |
| RSV-A | *Pneumoviridae* | | *-* | 100 | 0.026 | 0.333 | 0.135 | -0.285 | 0.230 | 0.892 | -74.660 | 1.9% | 0.524 | 1.067 | 1.349 | 0.643 |
| BVDV | *Flaviviridae* | | *+* | 123 | 0.178 | 0.456 | -0.226 | -0.311 | 0.377 | 0.901 | -48.109 | 0.7% | 0.788 | 0.929 | 0.950 | 0.661 |
| HPeV-3 | *Picornaviridae* | | *+* | 145 | 0.092 | 0.390 | -0.107 | -0.111 | 0.174 | 0.757 | -51.913 | 1.6% | 0.698 | 0.596 | 1.107 | 0.938 |
| CHIKV | *Togaviridae* | | *+* | 245 | 0.042 | 0.508 | -0.023 | -0.301 | 0.834 | 0.873 | -85.591 | 5.3% | 0.919 | 1.213 | 1.028 | 0.957 |
| EBOV | *Filoviridae* | | *-* | 200 | 0.001 | 0.413 | 0.086 | -0.154 | 0.591 | 0.730 | -68.068 | 2.2% | 0.840 | 0.890 | 1.353 | 1.118 |
| MeV | *Paramyxoviridae* | | *-* | 224 | 0.039 | 0.476 | 0.026 | -0.199 | 0.488 | 0.741 | -76.932 | -0.1% | 0.945 | 1.161 | 1.189 | 1.140 |
| EV-A71 | *Picornaviridae* | | *+* | 1161 | 0.148 | 0.464 | -0.208 | -0.325 | 0.524 | 0.777 | -63.715 | 0.7% | 0.725 | 1.295 | 0.822 | 1.421 |
| Porcine_KoV | *Picornaviridae* | | *+* | 90 | 0.121 | 0.520 | 0.537 | 0.341 | 0.640 | 0.570 | -84.142 | 16.9% | 0.925 | 1.796 | 0.852 | 1.469 |
| DENV1 | *Flaviviridae* | | *+* | 1556 | 0.066 | 0.464 | -0.208 | -0.325 | 0.424 | 0.583 | -59.423 | 1.8% | 0.745 | 1.542 | 0.935 | 1.575 |
| HEV | *Hepeviridae* | | *+* | 100 | 0.187 | 0.556 | 0.138 | 0.408 | 0.796 | 0.950 | -98.512 | 3.9% | 2.709 | 1.430 | 1.910 | 1.661 |
| HNoV_GGII | *Caliciviridae* | | *+* | 100 | 0.189 | 0.496 | -0.024 | -0.268 | 0.439 | 0.563 | -83.164 | 1.6% | 0.844 | 1.990 | 0.856 | 1.774 |
| OC43_gt2 | *Coronaviridae* | | *+* | 113 | 0.010 | 0.370 | -0.300 | 0.316 | 0.474 | 0.936 | -62.684 | 17.7% | 0.812 | 0.757 | 1.023 | 1.851 |
| JEV | *Flaviviridae* | | *+* | 62 | 0.088 | 0.516 | -0.193 | -0.236 | 0.603 | 0.531 | -69.491 | 1.4% | 0.971 | 2.156 | 0.924 | 1.886 |
| HPgV-1 | *Flaviviridae* | | *+* | 100 | 0.122 | 0.592 | -0.150 | 0.292 | 0.697 | 0.669 | -98.945 | 12.0% | 2.477 | 2.055 | 1.324 | 1.928 |
| MNV | *Caliciviridae* | | *+* | 63 | 0.102 | 0.569 | -0.017 | -0.009 | 0.621 | 0.487 | -85.137 | 7.1% | 1.539 | 2.587 | 1.191 | 2.079 |
| HCV-3a | *Flaviviridae* | | *+* | 820 | 0.085 | 0.557 | 0.033 | 0.090 | 0.715 | 0.825 | -83.319 | 8.7% | 1.372 | 2.978 | 0.998 | 2.110 |
| Canine_KoV | *Picornaviridae* | | *+* | 25 | 0.125 | 0.584 | 0.808 | 0.122 | 0.738 | 0.428 | -88.384 | 17.8% | 0.883 | 3.690 | 0.793 | 2.125 |
| HCV-2a | *Flaviviridae* | | *+* | 51 | 0.109 | 0.578 | 0.060 | 0.045 | 0.694 | 0.758 | -94.955 | 7.7% | 1.763 | 2.851 | 1.224 | 2.149 |
| HKU1 | *Coronaviridae* | | *+* | 27 | 0.002 | 0.320 | -0.316 | 0.446 | 0.453 | 0.958 | -50.191 | 9.6% | 1.250 | 0.692 | 1.841 | 2.163 |
| TGEV | *Coronaviridae* | | *+* | 38 | 0.022 | 0.375 | -0.188 | 0.126 | 0.474 | 0.818 | -57.014 | 8.8% | 0.548 | 1.091 | 0.782 | 2.265 |
| FMDV-O | *Picornaviridae* | | *+* | 246 | 0.106 | 0.536 | 0.094 | -0.166 | 0.815 | 0.437 | -77.621 | 11.7% | 1.059 | 3.095 | 1.029 | 2.266 |
| OC43 | *Coronaviridae* | | *+* | 178 | 0.008 | 0.366 | -0.290 | 0.305 | 0.466 | 0.925 | -62.069 | 17.5% | 1.543 | 1.034 | 1.992 | 2.503 |
| FMDV-A | *Picornaviridae* | | *+* | 97 | 0.112 | 0.536 | 0.094 | -0.151 | 0.819 | 0.478 | -88.156 | 12.1% | 1.015 | 3.118 | 0.997 | 2.559 |
| NL63 | *Coronaviridae* | | *+* | 61 | 0.009 | 0.345 | -0.275 | 0.490 | 0.413 | 0.876 | -50.191 | 8.6% | 1.144 | 0.945 | 1.510 | 2.654 |
| HCV-1b | *Flaviviridae* | | *+* | 102 | 0.094 | 0.587 | 0.060 | 0.044 | 0.736 | 0.765 | -88.349 | 8.5% | 1.810 | 3.991 | 1.238 | 2.905 |
| 229E_Camel | *Coronaviridae* | | *+* | 33 | 0.002 | 0.384 | -0.226 | 0.281 | 0.500 | 0.810 | -59.594 | 10.4% | 0.577 | 1.398 | 0.720 | 2.906 |
| 229E_Human | *Coronaviridae* | | *+* | 26 | 0.007 | 0.381 | -0.228 | 0.284 | 0.490 | 0.794 | -53.676 | 10.4% | 0.667 | 1.389 | 0.842 | 2.981 |
| MERS-CoV | *Coronaviridae* | | *+* | 26 | 0.005 | 0.412 | -0.033 | 0.244 | 0.559 | 0.897 | -59.594 | 15.7% | 1.104 | 1.840 | 1.386 | 2.986 |
| HCV-1a | *Flaviviridae* | | *+* | 355 | 0.083 | 0.587 | 0.070 | 0.065 | 0.720 | 0.777 | -88.349 | 9.0% | 2.094 | 4.734 | 1.441 | 3.019 |
| RUBV | *Matonaviridae* | | **+** | 73 | 0.0542% | 0.696 | 0.260 | 0.040 | 1.057 | 0.732 | -131.04 | 3.2% | 2.755 | 9.913 | 1.258 | 3.596 |
| SARS-CoV | *Coronaviridae* | | *+* | 22 | 0.000 | 0.408 | -0.038 | 0.082 | 0.465 | 0.800 | -61.506 | 13.5% | 0.667 | 2.714 | 0.911 | 4.177 |
| SARS-CoV-2 | *Coronaviridae* | | *+* | 17550 | 0.000 | 0.379 | -0.066 | 0.077 | 0.400 | 0.827 | -58.700 | 15.1% | 1.025 | 4.263 | 1.562 | 7.481 |

^1^Abbreviations: See Table 1

^2^MPD: mean pairwise uncorrected nucleotide distance

^3^Excess of the frequency of C over the frequency of G, or U over A

^4^Representation expressed as the observed frequency divided by expected frequency based on mononucleotide composition

^5^Ratio of G->A transitions to A->G transitions or C->U transitions to U->C transitions

^6^Corrected ratio based on nucleotide composition (see Results text).

FIGURE S1

UNROOTED PHYLOGENIES OF SEQUENCE ALIGNMENTS USED FOR HOMOPLASY ANALYSIS

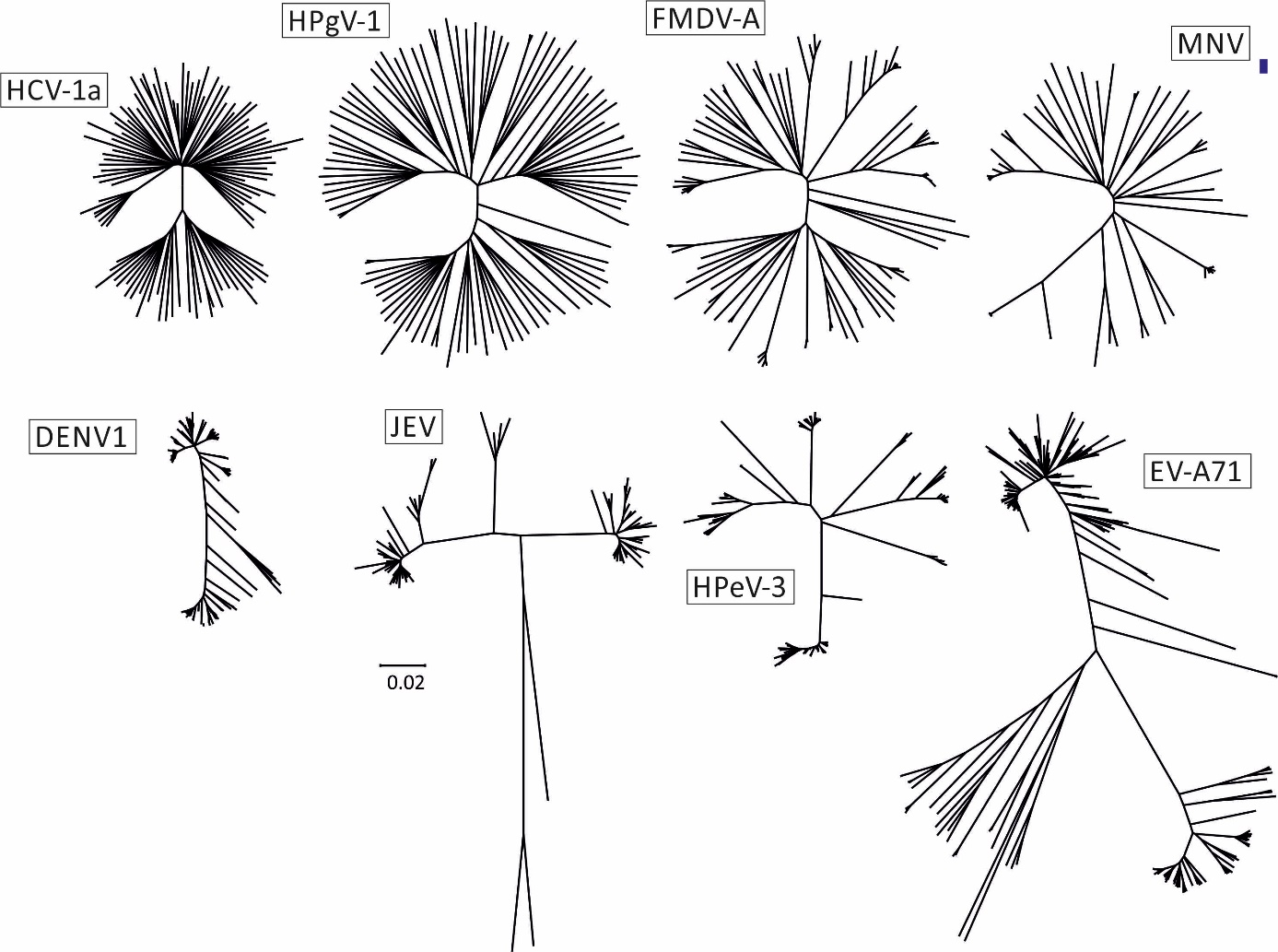

Neighbour-joining trees of coding region sequences used for lineage through time analysis plotted to the same scale (see scale). For comparability of the depicted trees, randomly selected subsets of 100 sequences were used for the larger HCV-1a, DENV1 and EV-A71 sequence alignments.

FIGURE S2

LIENEAGE THROUGH TIME PLOT FOR SEQUENCE ALIGNMENTS USED FOR HOMOPLAY ANALYSIS

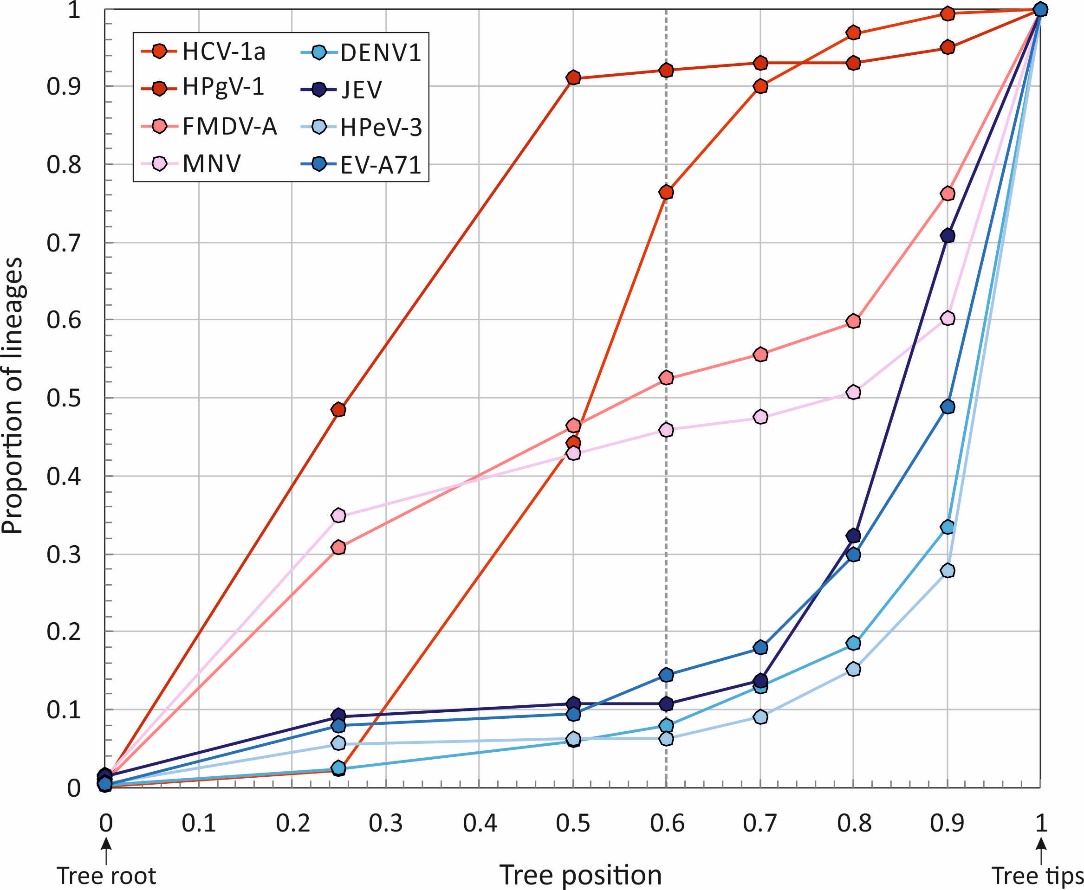

Proportion of total lineages (y-axis) at different positions in neighbour joining phylogenetic trees of virus alignments used for homoplasy analysis (x-axis). Virus groups were divided into those with high (red) and unbiased (blue) C->U / U->C transition asymmetries.
